# Rv0783c of *Mycobacterium tuberculosis* acts as a proton-motive force dependent multidrug efflux transporter involved in the efflux of structurally unrelated antibiotics and enhancing biofilm formation

**DOI:** 10.64898/2026.04.01.715825

**Authors:** Debleena Bhattacharyya, Debasmita Chatterjee, Aditya Prasad Panda, Anindya Sundar Ghosh

## Abstract

Despite multiple treatment strategies and extensive research on resistance mechanisms, tuberculosis (TB) remains a major global health threat, largely because of the rise of multidrug-resistant (MDR) and extensively drug-resistant (XDR) TB. Among various mechanisms complicating the situation, active antibiotic export via efflux pumps is particularly significant, yet largely unexplored. *Mycobacterium* sp. encodes numerous transporters, many of which are overexpressed in clinical isolates or under drug stress. Here, we examined the possible role of Rv0783c, a putative transporter that is reportedly overexpressed in drug-stressed conditions. Rv0783c conferred resistance to multiple structurally diverse antibiotics, fluoroquinolones and anti-TB drugs in the heterologous hosts, namely, *Escherichia coli* and *Mycobacterium smegmatis*. Reduced drug accumulation and active efflux of ethidium bromide (EtBr) confirmed its transport activity, which in turn gets nullified upon using the proton-motive force blocker, CCCP. On the other hand, its expression enhanced biofilm formation, linking antibiotic resistance to persistence-associated phenotype. Furthermore, site-directed mutagenesis confirmed the presence of crucial interacting residues with antibiotics that were identified by *in silico* analysis. Overall, we demonstrate the role of Rv0783c in the extrusion of first and second-line anti-TB drugs and enhancing biofilm formation.

## Introduction

In 2024, the global incidence of tuberculosis decreased for the first time since 2020, though it remained the primary cause of mortality from a single infectious agent, surpassing COVID-19 (Goletti *et al.,* 2025). Despite the progress in fighting tuberculosis, the WHO (World Health Organization) has reported that around 8.3 million new tuberculosis cases and 1.23 million fatalities still occur each year (World Health Organization, 2025). This highlights the need to strengthen the current knowledge and treatment strategies against the causative organism, *Mycobacterium tuberculosis* (*Mtb*) (World Health Organization, 2025). The emergence of MDR (multidrug resistant; resistant to at least isoniazid and rifampicin), XDR (extensively drug resistant; resistant to isoniazid, rifampicin, fluoroquinolones and one second-line anti-TB drug), and TDR (totally drug-resistant; resistant to all first and second-line anti-TB drugs) *Mtb* strains poses a significant challenge to the treatment regimen (Falzon, 2010).

Drug resistance in *Mtb* arises from both intrinsic and acquired mechanisms, including a highly impermeable mycolic acid-rich cell wall barrier, drug-modifying enzymes, and chromosomal mutations of drug target genes (Rattan et al., 1998). Interestingly, multiple drug-resistant *Mtb* clinical isolates lack mutations in known drug target genes, implying that alternative mechanisms may play a role in the development of resistance (Ramaswamy et al., 2003). Active drug efflux likely contributes to the resistant phenotypes lacking mutations (Machado et al., 2017). Efflux pumps (EP) are membrane-spanning proteins that extrude diverse substrates, including antimicrobials, toxins, heavy metals, metabolites, and quorum-sensing molecules, enhancing bacterial survival under stress (Huang et al., 2022). Although many efflux pumps in *Mycobacterium* species remain transcriptionally repressed under laboratory conditions, their activation plays a major role in multidrug resistance in clinical isolates (Huang et al., 2022). In addition to the overexpression of efflux pumps in response to prolonged drug exposure, several mutations in efflux pump genes have been reported in *Mtb* clinical and resistant isolates (AlMatar et al., 2019; Chimal-Muñoz et al., 2025). Genomic analysis reveals that putative drug transporters comprise about 10-12% of the total genome in *Mtb,* having one of the highest proportions of efflux pumps relative to genome size (Paulsen et al., 2001).

Among the broadly classified EP superfamilies, the primary active transporter ABC (<u>A</u>TP-<u>b</u>inding <u>c</u>assette) utilises ATP hydrolysis as an energy source, whereas secondary transporter families: MFS (<u>m</u>ajor facilitator <u>s</u>uperfamily), SMR (<u>s</u>mall <u>m</u>ultidrug resistance), RND (resistance-<u>n</u>odulation-<u>d</u>ivision), MATE (<u>m</u>ultidrug <u>a</u>nd toxic compound <u>e</u>xtrusion) superfamilies transporters are driven by proton motive force (PMF) (X. Z. Li & Nikaido, 2009). The MFS family is the largest among these secondary transporters (Pasqua et al., 2021). Most of the functionally studied efflux pumps belong to the MFS family, highlighting their contribution to multidrug resistance, not only in *Mtb* (De Rossi et al., 2002), but also in other bacterial species (Ghosh et al., 1998; Sigal et al., 2009). The differential prevalence of MFS subfamilies across *Mycobacterium* sp. reflects their possible role in adaptation in different niches. The pathogenic *Mtb* prioritises DHA (<u>D</u>rug-<u>H</u>^+^ <u>a</u>ntiporter subfamily, 58%) to survive antibiotics and host-immune stress, whereas the environmental species *Mycobacterium smegmatis* (*M. smegmatis*) prefers metabolite-H^+^ symporters (MHS) over DHA (25%) to acquire diverse nutrients (Singh & Akhter, 2025). Bioinformatics analysis using motif and sequence similarity has identified more than 20 putative MFS EPs in the organism, of which Tap (*rv1258c*), EfpA (*rv2846c*), JefA (*rv2459*), and P55 (*rv1410c*) are the major contributors (Aínsa et al., 1998; Gupta et al., 2010; Rai & Mehra, 2021; Santangelo et al., 2001). Prolonged exposure to sub-therapeutic concentrations of anti-mycobacterial drugs, often arising from poor TB treatment, can trigger efflux pump overexpression (Rindi, 2020). Several studies have reported that low-level resistance to fluoroquinolones (FQ), aminoglycosides, first-line anti-TB drugs, and even bedaquiline can arise from multiple EP overexpression or EP gene mutations, in the absence of target gene mutations. Few key contributors are *rv1258c*, *drrAB*, *pstB* and *rv2686c-2687c-2688c* for FQ-resistance (Pasca et al., 2004); *rv1258c* and *rv2333c* for aminoglycoside-resistance (Liu et al., 2019; Ramón-García et al., 2007a), *rv1877* for multidrug resistance (Adhikary, Biswal, & Ghosh, 2022) and mmpl5 (*rv0676)* for Bedaquiline resistance (Anthony Malinga & Stoltz, 2016).

Among many transporters linked to drug resistance*, rv0783c*, a putative MFS transporter, was reportedly overexpressed in multiple clinically resistant *Mtb* isolates (Bhattacharjee et al., 2022; G. Li et al., 2015; Machado et al., 2017). Multiple transcriptomic studies reported its enhanced expression in first-line anti-TB drug-resistant (isoniazid and rifampicin-resistant) (Pang et al., 2013) and second-line drug-resistant (amikacin and kanamycin) (Sowajassatakul et al., 2018) isolates. A few SNPs in the *rv0783c* locus are reported in rifampicin-resistant *Mtb* isolates lacking *rpoB* RRDR (<u>R</u>ifampicin resistant <u>d</u>etermining region) mutations (Jamieson et al., 2014). Although a steady-state basal level of *rv0783c* was observed in drug-susceptible *Mtb* isolates (Calgin et al., 2013a; Jiang et al., 2008), *rv0783c* remains functionally unexplored; its substrate specificity, transport mechanism, and physiological roles are yet to be elucidated. Considering the existing lacuna, our study explores the contribution of Rv0783c in exerting antimicrobial resistance (AMR). Additionally, we investigate whether its expression influences bacterial survivability through biofilm formation, which is associated with persistence and drug tolerance. Therefore, understanding the role of Rv0783c is essential to strengthen our knowledge on MFS transporter-mediated efflux in *Mtb*, and to evaluate its potential as a target for efflux inhibition strategies.

## Results

### Structural architecture and signature motifs support a transporter role of rv0783c

The gene *rv0783c* encodes a 56 kDa protein with 13 transmembrane helices (**Figure S1**). Multiple sequence alignment revealed the conserved MFS transporter signature motif A (GxLaDrxGrkxxxl, where x denoted any amino acid; capital and lowercase representing amino acid frequency of>70% and 40–70%, respectively) in between TM2-TM3 helix), motif B (lxxxRxxqGxgaa, in between TM3-TM4 helix), motif C or antiporter motif (gxxxGPxxGGxl, in TM5), which are functionally important for drug transporters (De Rossi et al., 2002)(**Figure S1**). The presence of 12 core transmembrane helices with one additional C-terminal helix places it closer to the DHA2 subfamily (14 helices) than that of the DHA1 subfamily (12 helices), supporting its classification as a DHA type MFS transporter (Reddy et al., 2012). In addition, Phyre2 Server predicted Rv0783c has 66% alpha-helix, 1% beta sheet and 33% loop region. Moreover, Phylogenetic tree analysis revealed that orthologs of Rv0783c are present in several *Mycobacterium* species, including pathogenic *M. bovis, M. marinum,* as well as NTMs (<u>N</u>on-tubercular *<u>M</u>ycobacteria,* e.g. *M. marinum, M. haemophilum, etc.*) (**Figure S1**). In contrast, *M. smegmatis* lacks an annotated ortholog. To further elucidate Rv0783c’s function *in vitro*, we have used *E. coli* and non-pathogenic, fast-growing *M. smegmatis* cells, which are routinely used as a model for *Mtb* (Lelovic et al., 2020).

### Rv0783c contributes to drug resistance in heterologous hosts

To investigate the role of Rv0783c in antibiotic resistance, we cloned the gene into inducible vectors pBAD18-Cam and pMIND and expressed it with 0.2% arabinose (for pBAD18-Cam in *E. coli*) and 20 ng mL^-1^ tetracycline (for pMIND in *M. smegmatis*). Protein expression was confirmed by membrane isolation and subsequently analysed by SDS-PAGE (**Figure S2**). We assessed the contribution of Rv0783c to multidrug resistance using the antibiotic susceptibility assay. The expression of *rv0783c* in *E. coli* led to 2-4-fold decrease in susceptibility values (as revealed by determining <u>M</u>inimum inhibitory <u>c</u>oncentrations, MICs) to multiple antibiotics, including, fluoroquinolones (sparfloxacin and lomefloxacin); aminoglycosides (kanamycin, neomycin, gentamicin, amikacin and apramycin); beta-lactams (ampicillin and oxacillin); and a 2-fold decrease in susceptibility for the dyes like ethidium bromide and Rhodamine B (**Table S1**). Similar variations were also noted in *M. smegmatis* cells. Expression of *rv0783c* in *M. smegmatis* also reduced the susceptibility to fluoroquinolones: sparfloxacin (4-fold), lomefloxacin (4-fold), norfloxacin and ciprofloxacin (2-fold); first line anti-TB drugs-rifampicin (8-fold), isoniazid (4-fold), ethambutol (2-fold); second-line anti-TB drug: amikacin (4-fold), and other antibiotics from different classes: apramycin (2-fold), gentamicin (2-fold) and ampicillin (4-fold) (**Table 1**).

**Table 1:** Comparative susceptibilities of *M. smegmatis* cells expressing pMIND control and Rv0783c towards different antimicrobials and compounds, in the absence and presence of sub-inhibitory concentration of CCCP.

| Antibiotic class | Antibiotics | Minimum Inhibitory Concentration (mg L<sup>-1</sup>) |  |  |  |
| --- | --- | --- | --- | --- | --- |
|  |  | Without CCCP |  | With CCCP |  |
|  |  | *M. smegmatis* | Rv0783c | *M. smegmatis* | Rv0783c |
| First-line anti-TB drugs | Rifampicin | 2 | 16 | 0.5 | 0.5 |
|  | Isoniazid | 32 | 128 | 4 | 4 |
|  | Ethambutol | 0.25 | 0.50 | 0.05 | 0.050 |
| Second-line anti-TB drugs (FQs) | Sparfloxacin | 2 | 8 | 0.12 | 0.12 |
|  | Lomefloxacin | 1 | 4 | 0.5 | 0.5 |
|  | Norfloxacin | 2 | 4 | 1 | 1 |
|  | Ciprofloxacin | 0.5 | 1 | 0.125 | 0.25 |
| Second-line anti-TB drugs (aminoglycosides) | Amikacin | 2 | 8 | 0.25 | 0.5 |
|  | Streptomycin | 1 | 4 | 0.25 | 0.5 |
| Other aminoglycosides | Apramycin | 2 | 4 | 1 | 2 |
|  | Gentamicin | 16 | 32 | 2 | 4 |
| Beta-lactams | Ampicillin | 64 | 256 | 8 | 8 |
|  | Oxacillin | 16 | 32 | 2 | 4 |
| Dye | Ethidium Bromide | 8 | 16 | 2 | 4 |
|  | Rhodamine B | 2 | 4 | 0.5 | 1 |

Rv0783c is predicted to encode a MFS transporter, relying upon the proton motive force (PMF) to mediate substrate transport (P. Li et al., 2017). To confirm whether its activity is PMF-dependent and the elevated resistance pattern of the host cells was due to *rv0783c* expression, we used the uncoupler CCCP (Carbonyl Cyanide m-Chlorophenylhydrazone) (Song & Wu, 2016) at a sub-inhibitory concentration (3 μg mL^-1^, 1/8^th^ MIC of CCCP). As expected, due to PMF disruption, both *E. coli* and *M. smegmatis* displayed increased drug susceptibility when treated with CCCP. For *M. smegmatis* cells expressing *rv0783c*, the resistance was reduced by 4 to 32-fold in the presence of CCCP, which was consistent with the results obtained in *E. coli* cells (**Table S1)**. Relative to the vector control, CCCP likely suppressed Rv0783c’s activity, as susceptibility values of sparfloxacin, ciprofloxacin, rifampicin and isoniazid were found similar between the control and test strains. For amikacin and ampicillin, the increased pattern of resistance persisted, though the fold change reduced to 2-fold as compared to the initial 4-fold **(Table 1**). The reduction in resistance profile indicates the inhibitory action of CCCP on the activity of Rv0783c, which might be due to the disruption of PMF. To determine whether this increased resistance was linked with the enhanced efflux activity, we next examined the ability of Rv0783c to extrude the fluorescent dye EtBr and other fluorescent antibiotics.

### Active efflux of EtBr by the cells expressing Rv0783c

Extrusion of fluorescent dyes (EtBr, Nile red) is often referred to as a gold standard for evaluating efflux activity (Blair & Piddock, 2016). The strong fluorescent signal of EtBr upon DNA binding allows the measurement of its intracellular concentration (Paixão et al., 2009). Cells were preloaded with dye in the presence of CCCP to equalise the initial fluorescence level for the control and test *M. smegmatis* cells. After energisation, cells expressing *rv0783c* demonstrated a rapid efflux compared to control cells carrying the control pMIND vector (**Figure 1a**). The t_efflux50%_ (time at which 50% of dyes efflux out from cells) (Iyer et al., 2015) for test strains was 30 s for EtBr, compared to 50 s (EtBr) for control *M. smegmatis* cells, providing the initial evidence that Rv0783c might act as an efflux pump.

**Figure 1:**
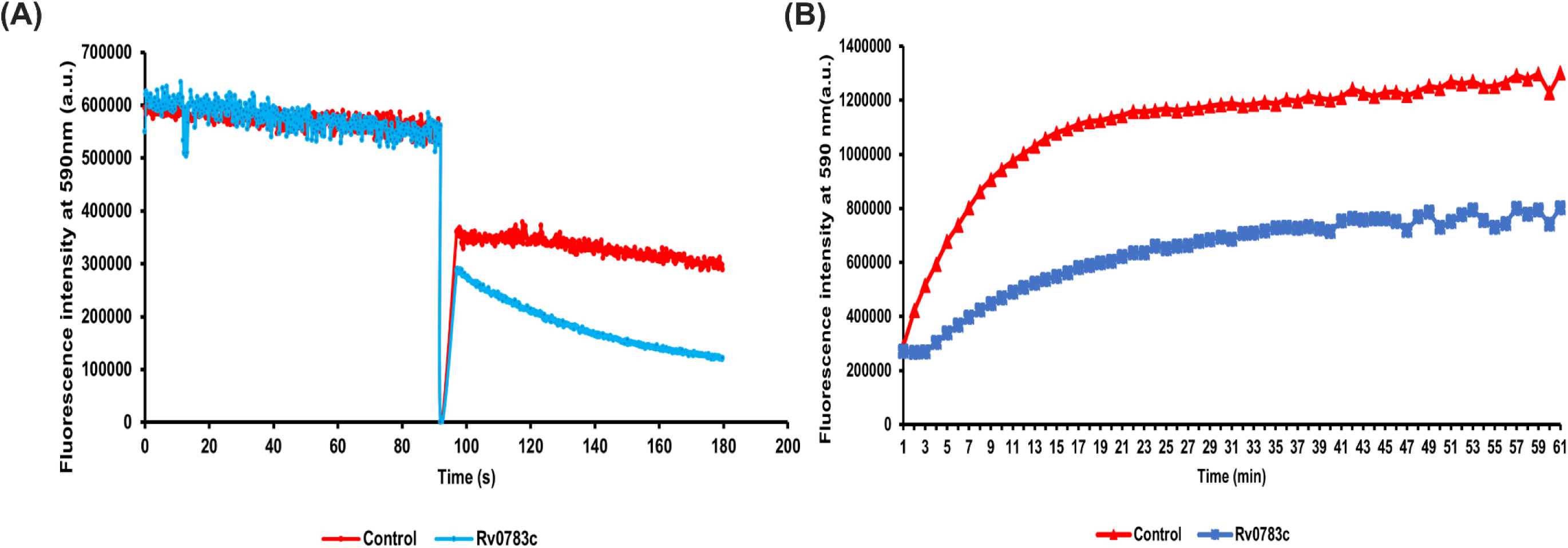
Real-time efflux and intracellular accumulation of EtBr in *M. smegmatis* cells expressing Rv0783c. (A) active efflux of EtBr from de-energised *M. smegmatis* cells expressing *rv0783c* or carrying the empty vector (control), following re-energisation by the addition of 50 mM glucose at 90 s. Fluorescence was monitored for a total duration of 180 s. Data represent the mean values from three independent experiments, with one set of representative data shown. (B) Intracellular accumulation of EtBr was measured at 590 nm at 1 min intervals for 60 min. Error bar represents mean ± standard deviation of three biological replicates.

### Decreased intracellular dye/drug level reflects Rv0783c-mediated efflux

To complement the results from the active efflux assay, we next performed time-dependent accumulation of EtBr and fluorescent antibiotics, i.e., sparfloxacin, lomefloxacin, norfloxacin, ciprofloxacin, and Bocilln FL with *E. coli* and *M. smegmatis* cells expressing *rv0783c* (Pu et al., 2016). The antibiotics and dyes were used at sub-inhibitory concentration (1/8^th^ of the MIC value), without compromising cell viability. We observed a notably lower accumulation of EtBr and antibiotics in cells expressing *rv0783c* as compared to the control cells. Within 10 min of EtBr exposure, *rv0783c*-expressing cells showed a reduction in EtBr accumulation than that of the vector control of *M. smegmatis* cells **(Figure 1b)**. Parallel assay in *E. coli* showed 57% reduction after 10 min of drug exposure, supporting its conserved efflux activity across different hosts. (**Figure S3**).

Accumulation of sparfloxacin was reduced by 45-50% in both hosts expressing *rv0783c* after 10 min of exposure **(Figure S3 and 2a)** as compared to the cells harbouring an empty vector. Similarly, comparable reductions were observed for the other fluoroquinolones tested. Following 10 min of antibiotic exposure, host cells expressing *rv0783c* showed lower levels of accumulation of lomefloxacin, norfloxacin, and ciprofloxacin by 49%, 48% and 43%, respectively **(Figure 3b, c, d)**. Likewise, for Bocillin-FL, a fluorescent penicillin analogue (G. Zhao et al., 1999), reduced accumulation (38%) after 5 min was observed than that of vector control *M. smegmatis* cells **(Figure 3e)**. Altogether, these findings suggest a general reduction in accumulation consistent with enhanced efflux **(Figures 2 and 3).** The addition of CCCP at 15 min led to a significant increase in fluorescence level, due to the increased accumulation in both the control and test strains, confirming the inhibitory action of CCCP on this efflux pump **(Figure S3 and 2).** Antibiotic accumulation by experimental cells was restored to the level of control cells upon CCCP treatment, reinforcing the previously observed reduction of MIC results in the presence of CCCP. Taken together, these results emphasise the role of Rv0783c in efflux-mediated multidrug resistance. To further determine the residues involved in the efflux activity of Rv0783c, we next conducted *in silico* analysis of interaction sites of antibiotics with the efflux pump.

**Figure 2:**
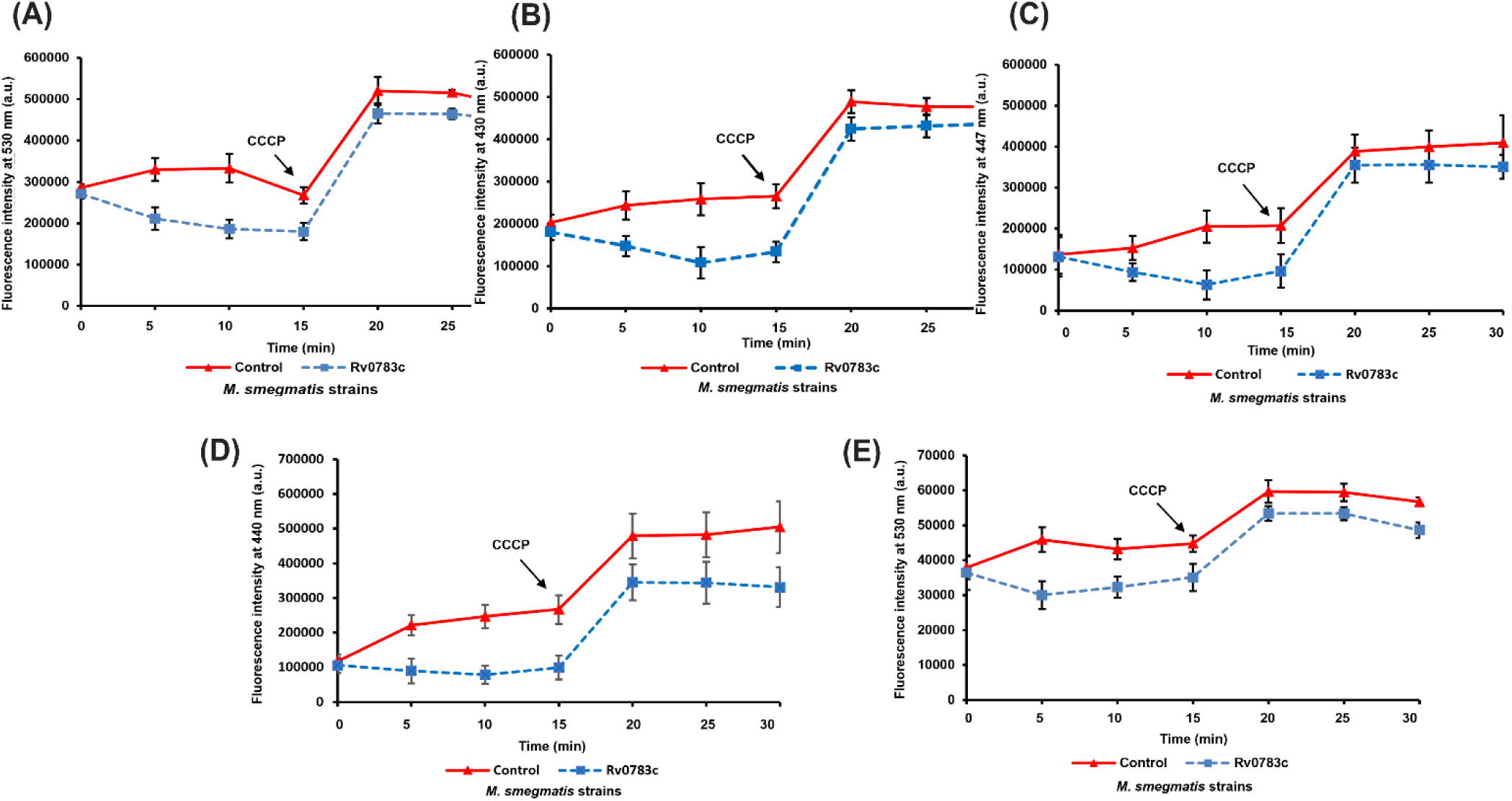
Intracellular accumulation of antibiotics in *M. smegmatis*. Accumulation of sparfloxacin (A), lomefloxacin (B), norfloxacin (C), ciprofloxacin (D), and Bocillin FL (E) at 5 min intervals for 30 min by cells expressing *rv0783c* (blue, closed square) compared to the vector control (red, closed triangle). Antibiotics and CCCP were added at 0 and 15 min, respectively. Error bar represents mean±standard deviation of three biological replicates (n=3).

**Figure 3:**
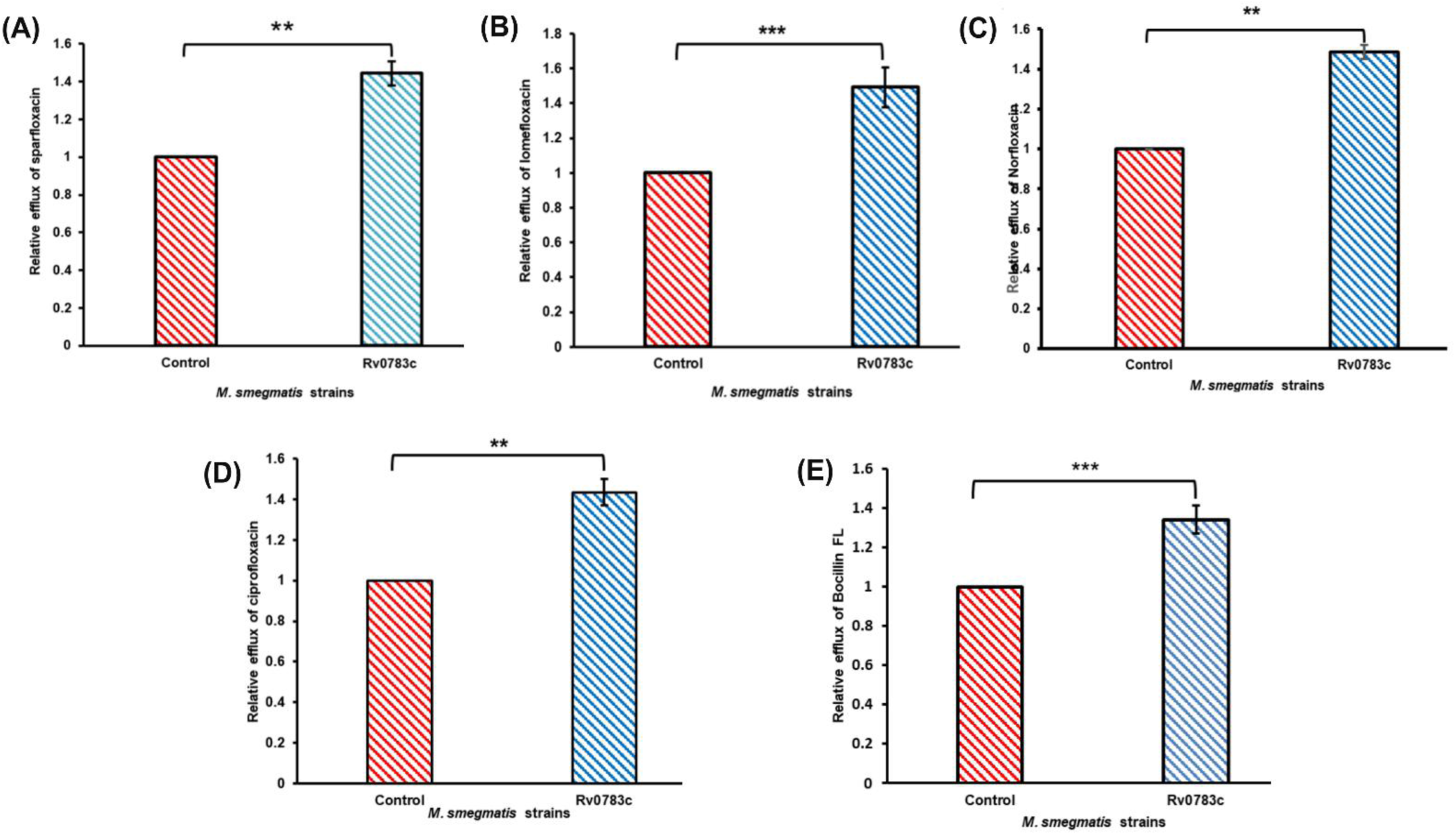
Relative efflux of antibiotics. Relative efflux values of sparfloxacin (A), lomefloxacin (B), norfloxacin (C), ciprofloxacin (D), and Bocillin FL (E) were calculated 10 min after antibiotic addition. Error bars indicate the mean ± standard deviation determined from replicates (n = 3). A two-tailed unpaired Student’s t-test was performed with the control and test datasets: ns: non-significant p>0.05, *p<.05, **p<0.01, ***p<0.001, ****p<.0001 The p-values were 0.003 (A), 0.001 (B), 0.001 (C), 0.002 (D),0.009 (E), respectively.

### In silico studies support the multidrug efflux activity of Rv0783c

Substrate docking analysis was performed to investigate the residue-level interactions of antibiotics. The 3D protein structure was retrieved from the AlphaFold database, and low-confidence N-terminal (1-37) residues were removed. Due to the absence of experimentally defined binding sites, blind docking was performed using a grid box over the entire protein. According to the “Goldilocks affinity” concept (Dey et al., 2020), unlike the enzyme-substrate complexes where high-affinity binding is observed, EPs rely on more transient interactions, since too tight binding might hinder transport (Drew et al., 2021). Findings from antibiotic susceptibility assays, with a 4-8-fold increase in resistance for sparfloxacin, isoniazid, and ampicillin, while a 2-fold change for norfloxacin and ciprofloxacin, were extrapolated and supported the binding affinity differences mentioned in **Table S2**. Most antibiotics docked along the central pore, with the key residues listed in **Table S2** and representative binding poses in **Figure 4**. Overall, the docking results aligned well with our experimental data, and accordingly, a few residues were subjected to site-directed mutagenesis to understand their contribution to the transport process.

**Figure 4:**
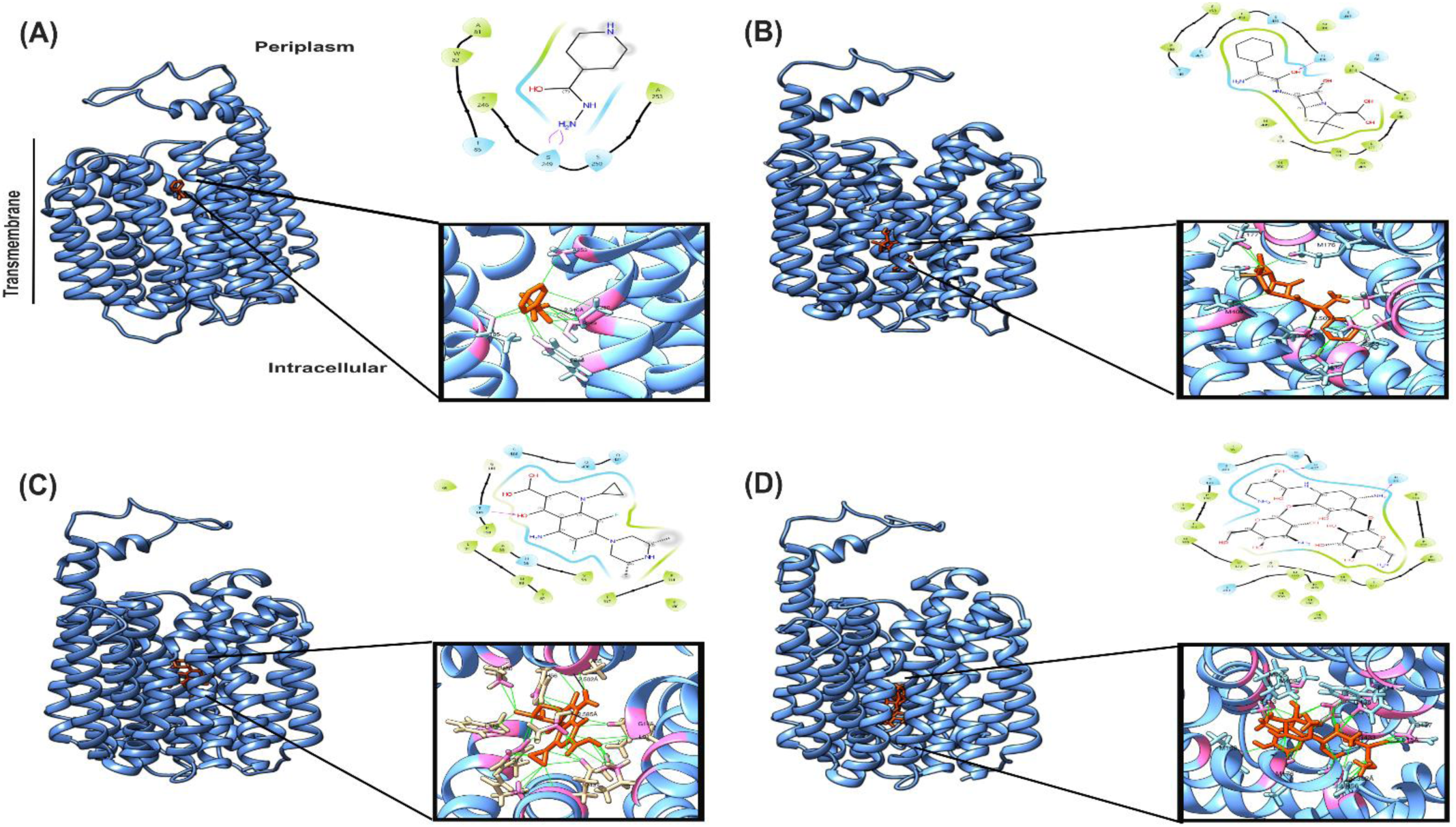
Molecular docking study of Rv0783c with different antibiotics: Interaction map with isoniazid (A), ampicillin (B), sparfloxacin (C), and amikacin (D) was performed using Autodock Vina and visualised by ChimeraX and Schrödinger Maestro. The broad channel view (left) and the corresponding interaction maps (right) were shown. The protein depicted in blue, antibiotics in orange, interacting residues in pink, bonds in black and favourable contacts in green.

### Substitutions of D58, R138, Q141 and H56 to alanine affect the efflux pump activity of Rv0783c

To identify the functional contributions of key residues of Rv0783c, we generated a series of site-directed mutants based on SNPs reported in clinical isolates and mechanistic insights of MFS pumps. According to the literature, two Rv0783c SNPs, G406V and F508S, have been identified in rifampicin-resistant isolates and clinical isolates from Vietnam, respectively (Hang et al., 2019; Jamieson et al., 2014). However, our functional analysis showed no increase in resistance pattern, suggesting that these two mutations alone might not be sufficient to alter the resistant phenotype (**Table S3 and Figure S4**). A review on mechanistic insights of MFS transporters by Drew *et al*., suggested the charged residues D^34^, R^112^ and Q^115^ are important for proton-coupled substrate translocation in MdfA and EmrD efflux pump (MFS family) (Drew et al., 2021). Guided by these findings, we performed sequence alignments with a few well-known MFS transporters to identify the corresponding positions in Rv0783c. Those charged residues (D^58^, R^138^, Q^141^), substituted with alanine separately, resulted in a loss-of-function phenotype. These single amino acid substitutions resulted in a similar susceptibility in terms of MIC as that of wild type (WT) *M. smegmatis* (**Table 2**). Likewise, increased accumulation of sparfloxacin and norfloxacin (**Figure 5**) was also observed compared to *rv0783c* expressing cells, supporting their predicted role in antibiotic efflux. Additional mutation sites selected from docking with antibiotics revealed that H56A might cause a substantial change in pump activity, whereas Q437A had no effect (**Table S3 and Figure S4**). Collectively, these results highlight the probable contribution of a few residues in the efflux activity.

**Figure 5:**
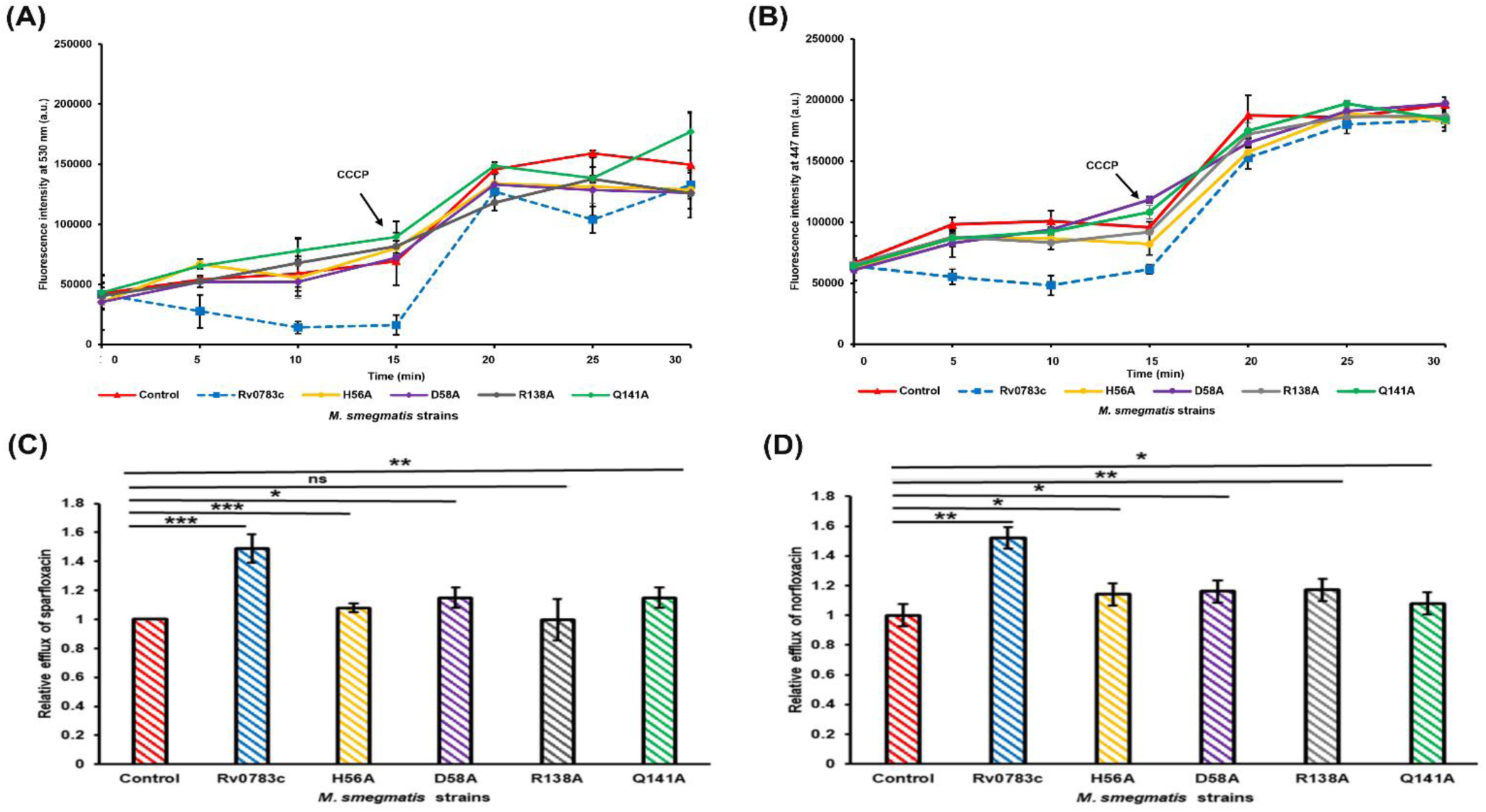
Comparative intracellular accumulation of antibiotics of wild-type and mutated Rv0783c. Accumulation of sparfloxacin (A), norfloxacin (B) at 5 min intervals for 30 min by cells expressing *rv0783c* (blue, closed square) compared to the vector control (red, closed triangle). Antibiotics and CCCP were added at 0^th^ and 15^th^ min, respectively. Relative efflux values of sparfloxacin (C), norfloxacin (D) were calculated 10 min after antibiotic addition. Error bars indicate the mean ± standard deviation determined from replicates (n=3). A two-tailed unpaired Student’s t-test was performed with the control and test datasets: ns: non-significant p>0.05, *p<.05, **p<0.01, ***p<0.001, ****p<.0001.

**Table 2:** Comparative susceptibilities of *M. smegmatis* cells expressing mutated Rv0783c, WT Rv0783c and vector control, towards different antimicrobials and compounds, in the absence and presence of sub-inhibitory concentration of CCCP.

| Antibiotic class | Antibiotics | Minimum Inhibitory Concentration (mg L<sup>-1</sup>) |  |  |  |  |  |  |  |  |  |  |  |
| --- | --- | --- | --- | --- | --- | --- | --- | --- | --- | --- | --- | --- | --- |
|  |  | Without CCCP |  |  |  |  |  | With CCCP |  |  |  |  |  |
|  |  | *M. smeg* | Rv0783c | H56A | D58A | R138A | Q141A | *M. smeg* | Rv0783c | H56A | D58A | R138A | Q141A |
| First-line anti-TB drugs | Rifampicin | 2 | 16 | 2 | 2 | 2 | 2 | 0.5 | 0.5 | 0.5 | 0.5 | 0.5 | 0.5 |
|  | Isoniazid | 32 | 128 | 64 | 32 | 32 | 32 | 4 | 4 | 4 | 4 | 4 | 4 |
|  | Ethambutol | 0.25 | 0.50 | 0.25 | 0.25 | 0.25 | 0.25 | 0.05 | 0.1 | 0.05 | 0.05 | 0.05 | 0.05 |
| Second-line anti-TB drugs (FQs) | Sparfloxacin | 2 | 8 | 2 | 2 | 2 | 2 | 0.12 | 0.12 | 0.12 | 0.12 | 0.12 | 0.12 |
|  | Lomefloxacin | 1 | 4 | 1 | 1 | 1 | 1 | 0.5 | 0.5 | 0.5 | 0.5 | 0.5 | 0.5 |
|  | Norfloxacin | 2 | 4 | 2 | 2 | 2 | 2 | 1 | 1 | 1 | 1 | 1 | 1 |
|  | Ciprofloxacin | 0.5 | 1 | 0.5 | 0.5 | 0.5 | 0.5 | 0.125 | 0.25 | 0.125 | 0.125 | 0.125 | 0.125 |
| Second-line anti-TB drugs (aminoglycosides) | Amikacin | 2 | 8 | 2 | 2 | 2 | 2 | 0.25 | 0.5 | 0.25 | 0.25 | 0.25 | 0.25 |
|  | Streptomycin | 1 | 4 | 1 | 1 | 1 | 1 | 0.25 | 0.5 | 0.25 | 0.25 | 0.25 | 0.25 |
| Other aminoglycosides | Apramycin | 2 | 4 | 2 | 2 | 2 | 2 | 1 | 2 | 1 | 1 | 1 | 1 |
|  | Gentamicin | 16 | 32 | 16 | 16 | 16 | 16 | 2 | 4 | 2 | 2 | 2 | 2 |
| Beta-lactams | Ampicillin | 64 | 256 | 64 | 64 | 64 | 64 | 8 | 16 | 8 | 8 | 8 | 8 |
|  | Oxacillin | 16 | 32 | 16 | 16 | 16 | 16 | 2 | 4 | 2 | 2 | 2 | 2 |
| Dye | Ethidium Bromide | 8 | 16 | 8 | 8 | 8 | 8 | 2 | 4 | 2 | 2 | 2 | 2 |
|  | Rhodamine B | 2 | 4 | 2 | 2 | 2 | 2 | 0.5 | 1 | 0.5 | 0.5 | 0.5 | 0.5 |

### Rv0783c expression increases biofilm formation

Apart from conferring resistance to antimicrobials, efflux pumps often play a role in bacterial physiology, virulence, and biofilm formation (Hajiagha & Kafil, 2023; Padilla et al., 2010), by transporting EPS (<u>E</u>xtracellular <u>p</u>olymeric <u>s</u>ubstances) components, quorum-sensing and inducer molecules to the extracellular environment (Alav et al., 2018). To study the effect of Rv0783c on biofilm formation ability in *E. coli* and *M. smegmatis*, we performed a semi-quantitative static biofilm formation assay. A significant change in biofilm formation was observed upon Rv0783c expression, with a 2-2.5-fold increase in both hosts. For *E. coli*, the biofilm formation index (BFI) increased from 0.18 in control cells to 0.38 upon *rv0783c* expression. A similar pattern was observed in *M. smegmatis*, where the BFI of control cells was 0.43, and *rv0783c* expression increased it to 0.98. The addition of CCCP (3 µg mL^-1^) led to a notable decrease in biofilm formation in both control and test strains, reestablishing its inhibitory action **(Figure 6a and b).** Similar to the efflux activity, the biofilm formation ability of the cells expressing mutated Rv0783c (Rv0783c-_H56A,_ Rv0783c-_D58A,_ Rv0783c-_R138A,_ Rv0783c-_Q141A_) were reduced as compared to the cells expressing wild-type Rv0783c as control **(Figure 6c).**

**Figure 6:**
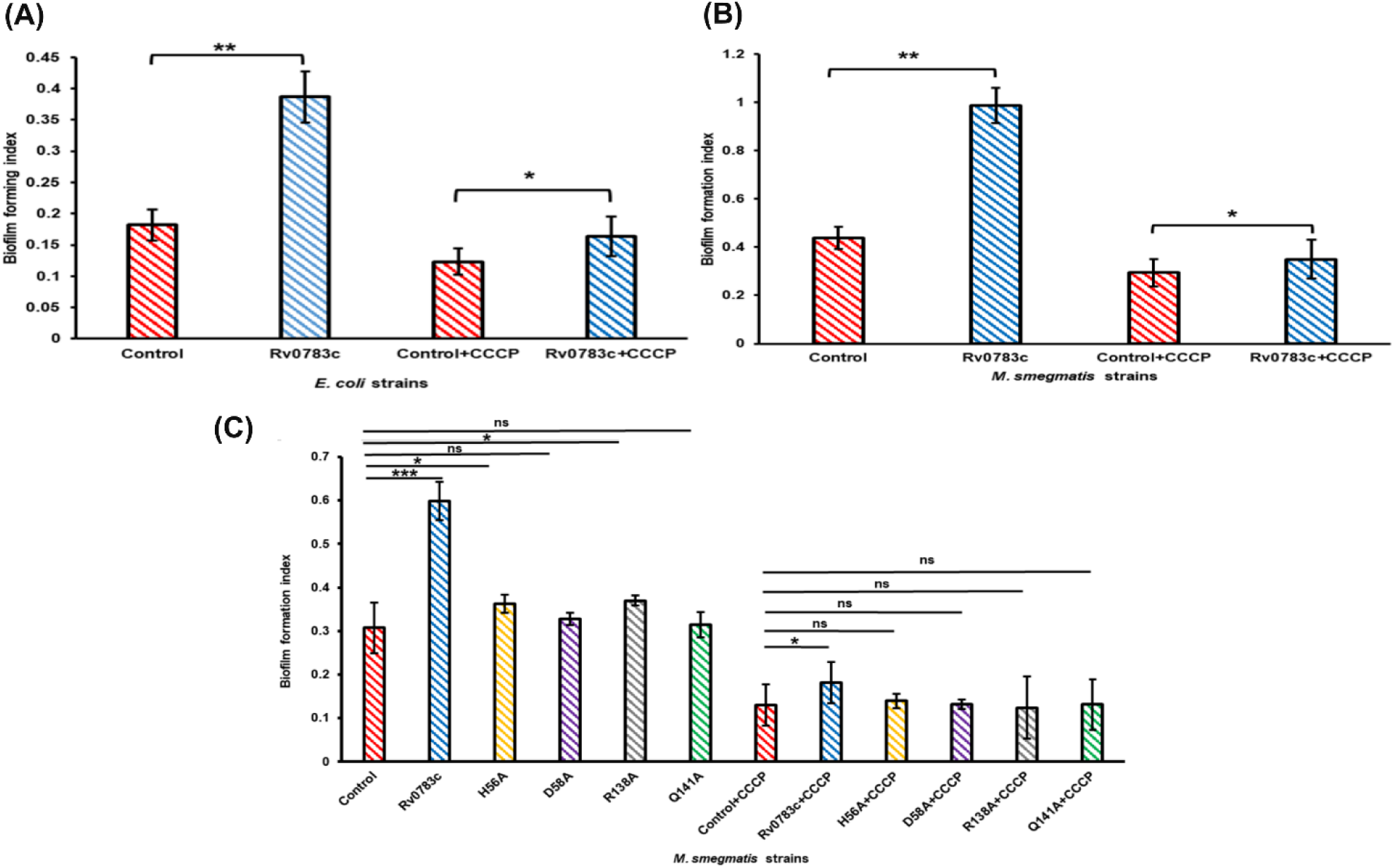
Semi-quantitative biofilm formation assay in *E. coli* and *M. smegmatis* with crystal violet. Biofilm formation by *E. coli* CS109 with pBAD vector (control) and Rv0783c (test) (A), *M. smegmatis* with pMIND vector (control) and Rv0783c (test) cells (B), respectively. All experiments were performed in quadruplicate and repeated three times. Statistical analysis was performed using two-tailed unpaired Student’s t-test; p>0.05, *p<.05, **p<0.01, ***p<0.001, ****p<.0001. Error bars indicate mean ± standard deviation (SD).

Microscopy data further validate this increase in biofilm formation. A denser biofilm structure was observed for wild-type Rv0783c-expressing cells compared to control *M. smegmatis* cells and mutated Rv0783c **(Figure S5).** Enhanced biofilm formation is often associated with alterations in phenotypic changes, including changes in colony morphology, pellicle formation and cell surface hydrophobicity (Limoli et al., 2015; Mirani et al., 2018).

Congo red (CR), a diazo dye, is reported to bind with (1-4) β-D-glucopyranosyl units of the cellulose matrix of mycobacterial biofilms(Bharti et al., 2021). Using both qualitative and quantitative assays, we found more CR deposition on the colony edges and 32% increase in CR binding upon *rv0783c* expression in *M. smegmatis* **(Figure S6a and b)**. Since mycobacterial species form thread-like pellicle structures on the air-liquid interface after extended incubation, we next examined whether this phenotype corresponded with the previous results. In support of our hypothesis, cells expressing *rv0783c* had slightly denser pellicle structure compared to the vector control **(Figure S6c)**.

We speculated that the overexpression of *rv0783c* might quantitatively alter extracellular polymeric substances (EPS) components. Using the chloroform: methanol extraction method, which separates the components into hydrophobicity-based phases, we observed a 2-fold increase in protein concentration and a 2-2.5-fold increase in carbohydrate content after expressing Rv0783c, whereas mutated versions (Rv0783c-_H56A,_ Rv0783c-_D58A,_ Rv0783c-_R138A,_ Rv0783c-_Q141A_) showed similar results as control *M. smegmatis* cells carrying empty vector (**Figure 7**). These results support the hypothesis that Rv0783c contributes to antibiotic persistence, perhaps through exporting antibiotics and molecules vital for biofilm formation.

**Figure 7:**
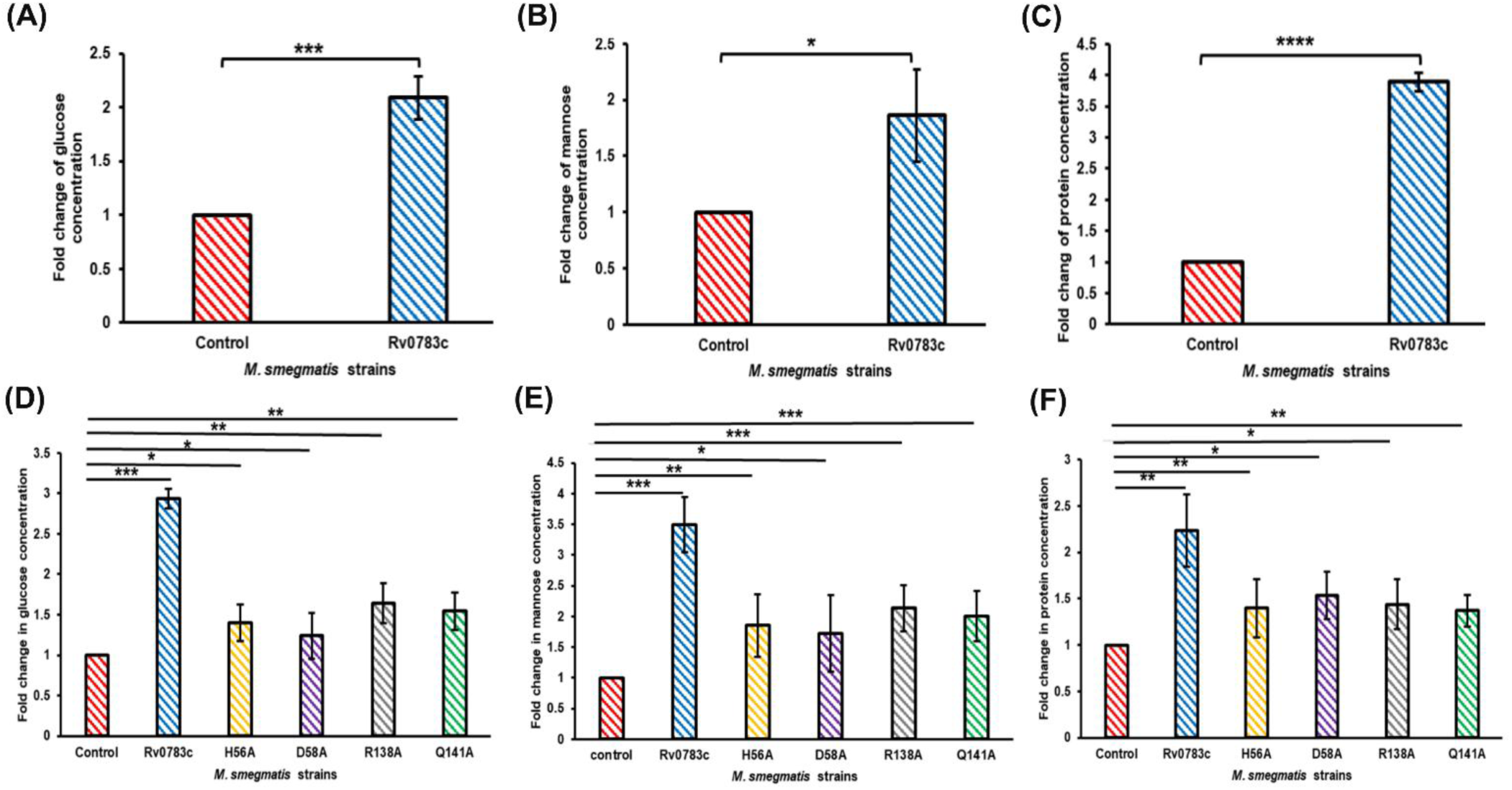
Altered ratios of EPS components in *M. smegmatis* cells expressing Rv0783c and its mutated versions. Fold change of carbohydrates: glucose (A), mannose (B) and protein (C) content in cells expressing WT *rv0783c* and mutated Rv0783c (D, E, F), respectively. Error bars indicate the mean ± standard deviation determined from replicates (n=3). A two-tailed unpaired Student’s t-test was performed with the control and test datasets: ns: non-significant p>0.05, *p<.05, **p<0.01, ***p<0.001, ****p<.0001.

## Discussion

Members of the MFS family efflux pumps, in particular the Drug/H^+^ antiporter (DHA) subfamily, function as proton-coupled transporters for a broad range of substrates, including metabolites, clinically significant drugs, and toxic compounds (Pao et al., 1998; Singh & Akhter, 2025). Though, earlier studies reported overexpression of *rv0783c* in multiple-drug-resistant *Mtb* isolates (Calgin et al., 2013b; G. Li et al., 2015), they lacked functional validation. In this context, the present study provides direct evidence of the protein’s importance in antibiotic resistance.

Conservation of Rv0783c across multiple *Mycobacterium* sp. (both tubercular and non-tubercular) suggests a possible role in survival and physiology. From the structure and sequence analysis, the presence of transmembrane helices and conserved MFS-DHA motifs provides the first clue of possible functional similarity to some well-studied MFS efflux pumps, such as NorA and QacA DHA pumps of *Staphylococcus aureus* (Brawley et al., 2021; Majumder et al., 2023). Heterologous expression of efflux pump genes has been widely used to directly evaluate their role in antibiotic resistance (Adhikary, Biswal, & Ghosh, 2022; Chatterjee, Daya Manasi, et al., 2024). Notably, consistent results have been found in both hosts, *M. smegmatis* and *E. coli,* suggesting that its activity is not specific to bacterial species. Upon ectopic expression of *rv0783c,* decreased susceptibility is observed not only to first and second-line anti-TB drugs but also to other classes of antibiotics, such as aminoglycosides, beta-lactams, and tetracyclines. Although Rv0783c mediates a low-to-moderate level of resistance, this phenomenon is clinically relevant as these drugs are a major backbone of the TB treatment regimen (Sotgiu et al., 2015). The observed behaviour naturally raises the question of how Rv0783c reduces susceptibility to such diverse compounds. Resistance to these drugs commonly involves mechanisms such as target modification, enzymatic drug inactivation and active efflux (Darby et al., 2022). The observed broad-spectrum resistance pattern suggests the later mechanism, a characteristic pattern of promiscuous MFS efflux pumps. For instance, MdfA of *E. coli* (Swick et al., 2011), EmrD-3 of *Vibrio cholerae* (Smith et al., 2009), and QacA of *S. aureus* (Mitchell et al., 1998), Rv1877 of *Mtb* (Adhikary et al., 2022), transport diverse antibiotics, antiseptics, and toxic dyes from cells.

The answer becomes clearer after the intracellular antibiotic accumulation assay and real-time efflux assay, which are considered gold-standard methods for drug efflux (Blair & Piddock, 2016). A persistent lower accumulation of the fluorescent dyes and antibiotics was observed in *rv0783c*-expressing cells over time. This inverse relationship between efflux activity and intracellular drug levels has been demonstrated in multiple reports, including the MSMEG_2991 pump of *M. smegmatis* (Bansal et al., 2016) and Rv2333c (stp) of *Mtb* (Ramón-García et al., 2007b). Passive diffusion and porin-mediated influx allow substrate entry (Prajapati et al., 2021) at comparable rates in both control and experimental sets, but the heterologous expression of efflux pump gene might prevent their intracellular build-up by rapid efflux, leading to a significant difference in accumulation profiles. Furthermore, disruption of PMF using an uncoupler like CCCP has been consistently used in studies to abolish efflux (Pule et al., 2016). CCCP acts by shuttling protons across the membrane in an uncontrolled manner, collapsing the proton gradient (Sekyere & Amoako, 2017), and inhibiting all proton gradient-dependent processes (Sanchez-Carbonel et al., 2021). Alongside increased antibiotic concentrations upon CCCP exposure, the differential antibiotic levels observed between the two sets were abolished, indicating that CCCP inhibits Rv0783c. This effect is further supported by a real-time efflux assay using EtBr (Paixão et al., 2009), in which, upon restoration of the proton gradient through glucose metabolism, a rapid, steeper decrease of fluorescence was observed in *rv0783c*-expressing cells compared to the control cells, suggesting active efflux. Together, these observations establish Rv0783c as a PMF-dependent MDR efflux pump.

*In silico* studies provided mechanistic insights into how Rv0783c accommodates such diverse antibiotics. MFS pumps possess a flexible central pore lined with multiple charged, hydrophobic and aromatic amino acids that can interact with structurally diverse substrates through H-bonds, hydrophobic, electrostatic and π-π interactions (Drew et al., 2021). Docking analysis revealed that all the antibiotics bind to the central channel region lined with charged residues (Majumder et al., 2019; Wu et al., 2020). Although the best-docked position differs for different classes, binding within the central pore suggests a common translocation pathway for structurally unrelated antibiotics. Previous work by Nie *et al*. suggests that due to the difficulty of substrate release during conformational change, substrates with higher binding affinity are less prone to export compared to substrates with lower binding affinity (Nie et al., 2016). Fitting well with the hypothesis, sparfloxacin, ampicillin, and isoniazid, which showed a higher fold change in resistance, have a higher binding free energy, implicating lower affinity; compared to other substrates like norfloxacin, which has a lower fold change in resistance and higher binding affinity. According to the rocker-switch model of substrate translocation, transitions between inward-open, occluded, and outward-open states are crucial (Sauve et al., 2023). Previous studies have shown the importance of protonation at the D^34^ residue for the transition to the occluded state in MFS pumps such as MdfA and QacA (Majumder et al., 2019; Sigal et al., 2009). In the MdfA efflux pump of *E. coli,* the R^112^ and Q^115^ residues of motif B are critical for the transition of occluded to inward-open state and proton coupling. Abolishment of these charged residues resulted in a loss of chloramphenicol resistance of MdfA (Y. Zhao et al., 2015). Although these residues are not conserved in Rv0783c, we could identify probable analogues through structure and sequence alignment analyses. Substituting those charged residues (D^58^, R^138^, Q^141^) with alanine likely disrupts the electrostatic interactions required for substrate translocation and conformational change, leading to the loss of efflux activity and biofilm formation. Docking with multiple antibiotics consistently identifies H^56^ as a key interacting residue within the central pore region. Previous study on the GlpT transporter of *E. coli* suggests the importance of a histidine residue (H^165^) of the central channel in stabilising the substrate through hydrogen and electrostatic bonds during transport (Sauve et al., 2023). Substituting H^56^ of Rv0783c to alanine likely abolishes these interactions, impairing substrate recognition, resulting in a loss-of-function activity. Based on the above findings, we can speculate the importance of these charged residues in substrate recognition and binding, and further translocation through the pump. The significance of these kinds of efflux pumps cannot be neglected, as they play a major role in primary resistance against a wide spectrum of antibiotics.

Efflux pumps have been increasingly reported as key contributors to biofilm formation beyond their role in antibiotic resistance (Vareschi et al., 2025; A. Zhao et al., 2023). Studies from diverse species like *Pseudomonas aeruginosa, Klebsiella pneumoniae, S. aureus, E. coli,* and *Mtb* have shown that this increase in biofilm formation can be linked with the extrusion of toxic metabolites, quorum-sensing molecules by these pumps, which maintain cellular homeostasis (McCarthy et al., 2015; Tuon et al., 2022). In the presence of efflux pump inhibitors (EPIs), active transport of intracellular metabolites and EPS-maturing components is abolished, resulting in a decrease in biofilm formation (Kvist et al., 2008; Reza et al., 2019; Sabatini et al., 2017). From our observations, the enhanced biofilm formation by *rv0783c* expression can be reversed by adding a PMF disruptor CCCP, which abolishes the active transport of toxic compounds. Another possible mechanism behind this increased biofilm formation upon efflux pump overexpression may be due to the increased transport of EPS (extracellular polymeric substance) components within the matrix (Hajiagha & Kafil, 2023). The increased protein and carbohydrate content in the EPS of cells expressing *rv0783c* supports this hypothesis.

Taken all together, *rv0783c* of *Mtb* encodes a multidrug efflux pump of the MFS family, conferring low to moderate level resistance to multiple antitubercular drugs and might play a role in efflux-mediated enhancement of biofilm formation. With its functional impact, it may represent a previously unexplored component of the efflux network of *Mtb*. From this study, we speculate few residues (H^56^, D^58^, R^138^, Q^141^) might play an important role in substrate translocation, though more structure-based validation could possibly be worthy to draw to further strengthen the conclusions.

## Material and Methods

### Bacterial strains, plasmids and culture conditions

*E. coli* XL1-Blue (*rec*A1 *end*A1 *gyr*A96 *thi*-1 *hsd*R17 *sup*E44 *rel*A1 *lac*) (Stratagene/Agilent, Santa Clara, CA, USA), *E. coli* CS109 (W1485 *glnV rpoS rph*)(Schnaitman & McDonald, 1984) used for cloning and preliminary assays, respectively, were cultured in Luria-Bertani Broth (LBB) or LB-agar (HiMedia, India) at 37 °C with proper antibiotics (12 µg mL^-1^ tetracycline for *E. coli* XL1B, 20 µg mL^-1^ chloramphenicol for strains having pBAD18-Cm vector(Guzman et al., 1995). *M. smegmatis* mc^2^155 (ATCC), used throughout the work, was grown in Middlebrook 7H9 broth (Hi-Media, Mumbai, MH, India) and 7H11 agar medium (Sigma-Aldrich, St. Louis, MO, USA) with oleic acid-ADC enrichment (Hi-Media Mumbai, MH, India), 0.05 % (w/v) Tween-80, 0.35 % (w/v) glycerol, and antibiotics (50 µg mL^-1^ kanamycin and 100 µg mL^-1^ hygromycin B for the strains having pMIND vector)(Williams et al., 2010). For antibiotic susceptibility testing of *E. coli* and *M. smegmatis*, Mueller-Hinton broth (MHB) (Hi-Media, Mumbai, MH, India) and 7H9 broth were used, respectively. Mycobacterial biofilms were cultured in 1X M63 medium (13.6 g KH_2_PO_4_, 2 g (NH_4_)_2_SO_4_, 0.5 mg FeSO_4_, 1 M MgSO_4,_ glucose (0.2%), distilled water up to 1,000 mL, and pH 7 to make 5X salt)(Polyudova et al., 2021). The polymerase and restriction endonucleases were obtained from New England Biolabs (Ipswich, MA, USA). Unless specified otherwise, all other reagents, including antibiotics and dyes, were purchased from Sigma-Aldrich (St. Louis, MO, USA).

### Construction of recombinant plasmids for in-vivo studies

For cloning, *rv0783c* was amplified from *M. tuberculosis* H37Rv gDNA with the specific set of primers mentioned in **Table S4**. The amplified fragment was cloned within *Nhe*I-*Hin*dIII sites in pBAD18-Cam (Guzman et al., 1995) and *Nde*I-*Spe*I sites in pMIND (Williams et al., 2010) to obtain pB0783c and pM0783c recombinant plasmids. The clones were confirmed by sequencing Eurofins Scientific (Hyderabad, TS, India). *E. coli* XL1-Blue was the cloning host for both constructs. Chemically competent *E. coli* CS109 cells were transformed with pB0783c plasmid (by heat-shock), and *M. smegmatis* cells were transformed with pM0783c plasmid by electroporation. For expression of the protein in *E. coli* CS109 and *M. smegmatis,* arabinose (0.2%)(Nelson & Young, 2000) and tetracycline (20 ng mL^-1^) were used, respectively (Pandey et al., 2018).

### Structure analysis and phylogenetic preparation

The protein sequence was obtained from Mycobrowser (Kapopoulou et al., 2011) and aligned with other MFS transporters of *M. tuberculosis* using Clustal-omega software(Sievers & Higgins, 2018) to identify the signature motifs of the MFS family in the sequence. The protter web server (Omasits et al., 2014) was used to predict the transmembrane helices, which are indicative of membrane-embedded transporter proteins. 3D structure was retrieved from the AlphaFold database(Varadi et al., 2022). Homologous sequences of Rv0783c were collected from the NCBI protein database using BLASTp with an e-value cutoff of 0.00001 and a query coverage of 100%. A phylogenetic tree was constructed in MEGA12 (Kumar et al., 2024) using the Maximum Likelihood method (Jones-Taylor-Thornton (1992) model of amino acid substitution). Node support was evaluated with 1,000 bootstrap replicates, and the tree with the maximum log likelihood value (−7,379.78) was selected.

### Ectopic expression of rv0783c in E. coli and M. smegmatis cells

Overnight-grown cultures of *E. coli* and *M. smegmatis* were inoculated (0.1%) into 10 mL of LBB and 7H9B, respectively, and grown at 37 °C, 150 rpm until early-log phase. Cultures were induced with arabinose (0.05-0.2%, w/v) for *E. coli* and tetracycline (20 ng mL^-1^) for *M. smegmatis* cells, for 14-16 h. Cells were harvested at 8,000 x g, 5 min (*E. coli*)/20 min (*M. smegmatis*) at 4 °C in an Eppendorf 5810R centrifuge (Hamburg, Germany) and washed twice with 1 mL of 10 mM Tris-Cl buffer (pH 7). Protease inhibitor PMSF (Phenylmethylsulfonyl fluoride) was added (1 mM) and sonicated, with 15 s on/off cycle, for 45 s (*E. coli*) and 60 s (*M. smegmatis*) on ice, followed by centrifugation at 50,000 x g for 5 min at 4 °C using Beckman Coulter Avanti JXN-26 ultracentrifuge (Brea, California, USA). The collected supernatants were subsequently centrifuged at 50,000 x g for 90 min at 4 °C. The pellet fractions were resuspended in 10 mM Tris-Cl buffer and treated with 2% sarkosyl (w/v) (sodium lauroyl sarcosinate) at 37 °C, shaking at 300 rpm in a thermomixer (Eppendorf, Hamburg, Germany) to solubilise the membrane. After a final centrifugation at 50,000 x g for 90 min at 4 °C, supernatant fractions were collected for protein estimation by Bradford assay, and finally, the samples were visualised after running through 12% SDS-PAGE (Sodium dodecyl sulfate-polyacrylamide gel electrophoresis) (Chatterjee, Panda, et al., 2024).

### Antibiotic susceptibility assay with different classes of antibiotics and dyes

Antimicrobial susceptibility of cells harboring *rv0783c* was measured by determining the minimum inhibitory concentration (MIC) of different classes of antibiotics namely, fluoroquinolones (norfloxacin, ofloxacin, sparfloxacin, ciprofloxacin, levofloxacin, moxifloxacin), aminoglycosides (apramycin, amikacin, gentamicin, kanamycin, neomycin), tetracycline, doxycycline, anti-TB drugs (rifampicin, isoniazid, ethambutol), dyes (ethidium bromide, rhodamine B) following CLSI guidelines(Lewis et al., 2023). MICs were determined using the micro-broth dilution assay method in a 96-well plate with serial dilutions of antibiotics, with an assay volume of 300 µL using MH broth (for *E. coli*) or 200 µL with 7H9 broth (for *M. smegmatis*), inoculated with 10^5^ cells per well(Adhikary, et al., 2022). Arabinose (0.2%, w/v) and tetracycline (20 ng ml^-1^) were used for inducing the expression, respectively, for *E. coli* and *M. smegmatis*. CCCP was used at a sub-inhibitory concentration (3 µg mL^-1^). After incubating at 37 °C for 14-16 h (*E. coli)* and 48-72 h (*M. smegmatis*), the optical density was measured at 600 nm with iMark^TM^ micro-plate reader (Bio-Rad, California, United States). All the experiments were performed in triplicate with six repeats to confirm reproducibility.

### Real-time efflux assay with Ethidium bromide

EtBr efflux of *M. smegmatis* cells was studied in real time with some minor changes (Adhikary et al., 2023). *M. smegmatis* culture was grown in 7H9 broth at 37 °C for 14–16 h after inducing at an OD_600_ of ∼0.4-0.6 with 20 ng mL^-1^ tetracycline for protein expression. Culture was harvested at 4,000 x g, 20 min, RT; using an Eppendorf 5810R (Hamburg, Germany) centrifuge and washed twice with 50 mM PPB supplemented with 1 mM MgCl_2_ (PPB), resuspended in the same buffer to adjust final OD_600_ of 1. All the subsequent steps were performed in RT. Following a resting period of 15 min, 2 mL aliquots were taken in 5 mL tubes. Cells were treated with CCCP (3 µg mL^-1^) and incubated for 15 min. EtBr (1.5 μM) was then added, kept at a 37 °C incubator with shaking at 150 rpm for 4 h. After an additional 1 h resting period, cells were pelleted by centrifugation, and the supernatant was completely removed. The pellet was resuspended in PPB, diluted tenfold, and fluorescence measurements were immediately taken for 90 s (at excitation-emission wavelength at 530 nm and 590 nm) using SpectraMax iD5 Microplate Reader (Molecular devices, San Jose, CA, USA). Efflux assessment was initiated by energising the cells with 50 mM glucose, and fluorescence was recorded further for 100 s. The experiment was done in duplicate and repeated thrice, with the representative results presented.

### Ethidium Bromide accumulation assay

This experiment was carried out with slight modifications to the previously described protocol by Rodrigues *et al*.(Rodrigues et al., 2011). *M. smegmatis* cells, carrying pM0783c, were washed with 50 mM potassium phosphate buffer (PPB with 0.05% Tween-80 and 1 mM MgCl_2_). The OD_600_ was adjusted with PPB to 0.4, and glucose (0.4%) was added to energise the cells. Ethidium bromide (EtBr) was added (final concentration 1.5 µM), and accumulation was recorded for 1 h, at every 1 min interval using SpectraMax iD5 Microplate Reader (Molecular devices, San Jose, CA, USA) with excitation at 530 nm and emission at 590 nm. The experiment was repeated thrice, each with technical duplicates to ensure consistency.

### Intracellular antibiotic accumulation assay

Accumulation assays for fluoroquinolone antibiotics (sparfloxacin, lomefloxacin, norfloxacin, ciprofloxacin) and Bocillin FL were performed based on previously established protocols (Chatterjee, Panda, et al., 2024; Ghosh et al., 1998) with a few minor modifications. *E. coli* and *M. smegmatis* cells were induced with 0.2% arabinose and 20 ng mL^-1^ tetracycline, respectively. Cells were washed with 50 mM PPB (pH 7) thrice, optimised at OD_600_ value of 1.0, and 0.4% glucose was added to energise the cells for 30 min at 37 °C. Antibiotics (sparfloxacin, lomefloxacin, ciprofloxacin, norfloxacin, Bocillin FL) were added to the suspension at sub-inhibitory concentrations (10-20 mg L^-1^), and aliquots (500 µL) were withdrawn at 5 min intervals for 30 min. CCCP (3 µg mL^-1^) was added at 15 min, and further aliquots were withdrawn. Cells were immediately washed thrice with PPB and resuspended in 0.1 M Glycine-HCl (pH 3) and incubated for 1 h (*E. coli*) or 14-16 h (*M. smegmatis*). After centrifugation, fluorescence intensity of supernatants was measured with a spectrofluorometer (FluoroMax 4; HORIBA Scientific Instruments, Japan) at an excitation-emission wavelength of 262 nm & 530 nm for sparfloxacin, 280 nm & 430 nm for lomefloxacin, 275 nm & 440 nm for ciprofloxacin, 281 nm & 447 nm for norfloxacin, 488 nm & 530 nm for Bocillin FL. The relative efflux of antibiotics was calculated using the following formula: Relative efflux (RE)= 1+(N_control_-N_test_)/N_control_, where N_control_ represents the antibiotic accumulated by control cells harbouring empty vectors, N_test_ represents the antibiotic accumulated by test strains. The formula for the relative percentage of efflux: (N_control_-N_test_/N_control_ X 100) (Adhikary, Biswal, Chatterjee, et al., 2022). All experiments were repeated three times in technical duplicates for consistency.

### In silico docking analysis

Structural modelling and molecular docking studies of Rv0783c were carried out to elucidate its structural motifs and substrate binding interactions. Due to the absence of a resolved crystal structure, the predicted 3D model of Rv0783c was retrieved from the Alphafold database (Varadi et al., 2022). The quality and reliability of the model were further validated with the Procheck tool. The putative binding sites and channel region were predicted using the Dogsite Scorer server (Volkamer et al., 2012). The protein structure was prepared for docking by energy minimisation using the Amber14ff force field, applying 500 steps of conjugate gradient and 100 steps of steepest gradient. Subsequently, ligand protonation was carried out, Gasteiger charges were assigned, and the files were saved in PDB format. The grid box is made at 77.024, 76.959 and 77.124 of x centre, y centre and z centre, respectively, for the target protein. Different classes of antibiotics were docked with AutoDock-Vina, and the results were visualised using UCSF-Chimera (Pettersen et al., 2004). The antibiotic structures were downloaded from the PubChem database (Kim et al., 2025). High-resolution visualisation representations of protein-ligand interaction were created with Schrödinger Maestro (*Schrödinger Release Notes - Release 2025-2*, n.d.).

### Site-Directed Mutagenesis

The plasmid pB0783c was used as a template for site-directed mutagenesis. With specific primer pairs (**Table S1**) and *Pfu* Turbo Polymerase (Agilent Technologies, Santa Clara, CA, USA) (Adhikary, Biswal, Chatterjee, et al., 2022), seven amino acid substitutions were carried out. Amplicons were treated with *Dpn*I and transformed into chemically competent *E. coli* XL1-Blue cells. All the mutations were confirmed by commercial sequencing by Eurofins Scientific (Hyderabad, TS, India). pB0783c plasmids, having specific mutations, were cloned separately in the pMIND vector and transformed into *M. smegmatis* cells.

#### Semi-quantitative assay for biofilm formation

Static biofilm formation assay was performed in a 24-well polystyrene plate as described, with a few changes (Bansal et al., 2016; De Siena et al., 2020). The well-grown primary cultures of *E. coli* and *M. smegmatis* strains were washed twice with PBS, diluted to an OD_600_ of 0.6 and 0.006, respectively, and inoculated in the wells of a 24-well plate having 1 mL of one-fifth LBB (*E. coli*) or 1X M63 salt (*M. smegmatis*). Plates were incubated for 48 h (*E. coli*)/7 days (*M. smegmatis*) at 37 °C in the static incubator. Following biofilm formation, the contents of the wells were aspirated out carefully and washed three times with PBS to remove non-adherent planktonic cells. Each well was treated with 1 mL of a 0.1% crystal violet solution (w/v) for 15 min. The stain was removed, washed with sterile water, and air-dried. Acetic acid (1 ml of 33% v/v) was added to each stained well, mixed by pipetting, and 200 μL transferred to a 96-well plate to measure the OD_600_ using a plate reader (Bio-Rad iMark^TM^microplate reader). To check the activity of inhibitor CCCP, the same experiment was repeated with media having CCCP at a sub-inhibitory concentration (3 μg mL^-1^). The Biofilm formation index (BFI) was calculated using the formula: BFI (OD_600_ of acetic acid solubilised stained wells-OD_600_ of control wells)/OD_600_ of planktonic cells. The formula for calculating biofilm inhibition percentage = [A_control_- A_treated_)/A_control_] X 100.

### Bright-field microscopic analysis of biofilm

Qualitative assessment of biofilm formation was conducted using bright-field microscopy. Biofilms were developed on microscopic cover slips placed in the wells of a 24-well polystyrene plate with 1 mL of 1X M63 media inoculated with 10^5^ *M. smegmatis* cells and incubated at 37 °C for 5-6 days. The cover slips were removed after incubation, washed with 1X PBS, stained with 0.1% CV(w/v) and observed using OLYMPUS 1×71 (Olympus Corporation, Tokyo, Japan) microscope(Pal et al., 2019).

### Congo-Red binding assay and Pellicle formation assay

Mycobacterial cultures were pelleted at 4,100 x g for 20 min, washed with sterile PBS, and 5 µL of each resuspended culture (OD_600_ ∼0.06) was spotted on 7H11 plates containing 100 µg mL^-1^ hygromycin, 40 µg mL^-1^ Congo red (CR), and 20 µg mL^-1^ Coomassie Brilliant Blue (CBB). Plates were kept at a 37 °C static incubator for 3 days to observe the CR binding(Pandey et al., 2018). To confirm further, the cultures were grown to the stationary phase in 7H9 broth supplemented with Congo red at 100 µg mL^-1^ and 0.05% Tween 80. The cells were subsequently rinsed thoroughly with water until the supernatants became colourless. The Congo red dye linked to the cells was removed using 400 μL of acetone for 1 h with moderate agitation. The absorbance of the extract was quantified at 488 nm. The Congo red binding index was determined as the absorbance at 488 nm of the acetone extracts divided by the optical density at 600 nm of the culture (Petchiappan et al., 2020). To facilitate pellicle formation, cultures comprising approximately 10^5 cells were cultivated in 10 mL of M63 media (lacking Tween 80) in sterile glass tubes at 37 °C without shaking, with results examined one-week post-inoculation (Pandey et al., 2018).

### Isolation of EPS from mycobacterial biofilm

Extraction of extracellular polysaccharides was carried out using the method described by Gopinath et al(Gopinath et al., 2024). Briefly, *M. smegmatis* cells were inoculated into 5 mL of 1X M63 media to form a biofilm. Biofilm-forming cells adhering to the glass tubes were scraped and inoculated into 10 mL of fresh 1X M63 media (without Tween 80) and incubated statically at 37 °C until a thick biofilm developed. The media was carefully removed and washed with 1X PBS. Using chloroform: methanol (2:1), adhered biofilm was collected and centrifuged at 5,000 rpm for 3 min to separate the hydrophilic, hydrophobic and cell phases. The upper methanol phase and lower chloroform phase contained hydrophilic and hydrophobic components, respectively, while the interphase consisted of the cellular pellet. Quantitative estimation of polysaccharides was performed using the phenol-sulfuric acid method, as modified for a 96-well plate format(Masuko et al., 2005). Glucose and mannose solutions (0-100 µg mL^-1^) were used as standard solutions to obtain the standard curve. To 50 µL of the upper hydrophilic phase, 50 µL of 5% phenol was added, followed by 150 µL conc. H_2_SO_4_. The plate was kept on the shaker for 10 min. Absorbance at 480 nm and 490 nm was measured for the quantification of pentose (mannose) and hexose (glucose) sugars, respectively, using SpectraMax iD5 Microplate Reader (Molecular devices, San Jose, CA, USA). Protein quantification of EPS was performed using the Bradford method, adapted for 96-well plate format (Filgueiras & Borges, 2022). 20 µL of the upper hydrophilic layer was taken in a 96-well plate, followed by the addition of 100 µL of Bradford reagent (SRL, Mumbai, MH, India). The solutions were mixed and incubated in the dark for 10 min at RT. Absorbance was measured at 595 nm using an iMark^TM^ microplate reader (Bio-Rad, California, United States). A standard curve was prepared using bovine serum albumin (BSA, concentration range: 0-100 µg mL^-1^).

## Statistical analysis

All experiments related to the accumulation of antibiotics and dyes, active efflux, and biofilm formation were performed in duplicate and repeated three times. Results were calculated as mean±standard deviation. The statistical significance (p-value) was calculated using GraphPad by performing two-sample, unpaired Student’s t-tests where ns = non-significant p-value > 0.05, *p<0.05, **p<0.01, ***p<0.001 and ****p<0.001.

## Author contributions

DB performed the experiments, analysed the data and wrote the manuscript draft. DC contributed to data curation and analysis, manuscript preparation. APP contributed to *in silico* works, analysis and manuscript preparation. ASG provided overall supervision, conceptualisation, experiment design and manuscript editing. All authors read and approved the final manuscript.

## Acknowledgement

The authors are thankful to Prof. Manikuntala Kundu, Bose Institute, Kolkata, for the genomic DNA of *M. tuberculosis* H37Rv.

## Data availability statement

All data generated or analysed during this study are included in this article and its supplementary section.

