## Supplementary data for "Rv0783c of *Mycobacterium tuberculosis* acts as a proton-motive force dependent multidrug efflux transporter involved in the efflux of structurally unrelated antibiotics and enhancing biofilm formation"

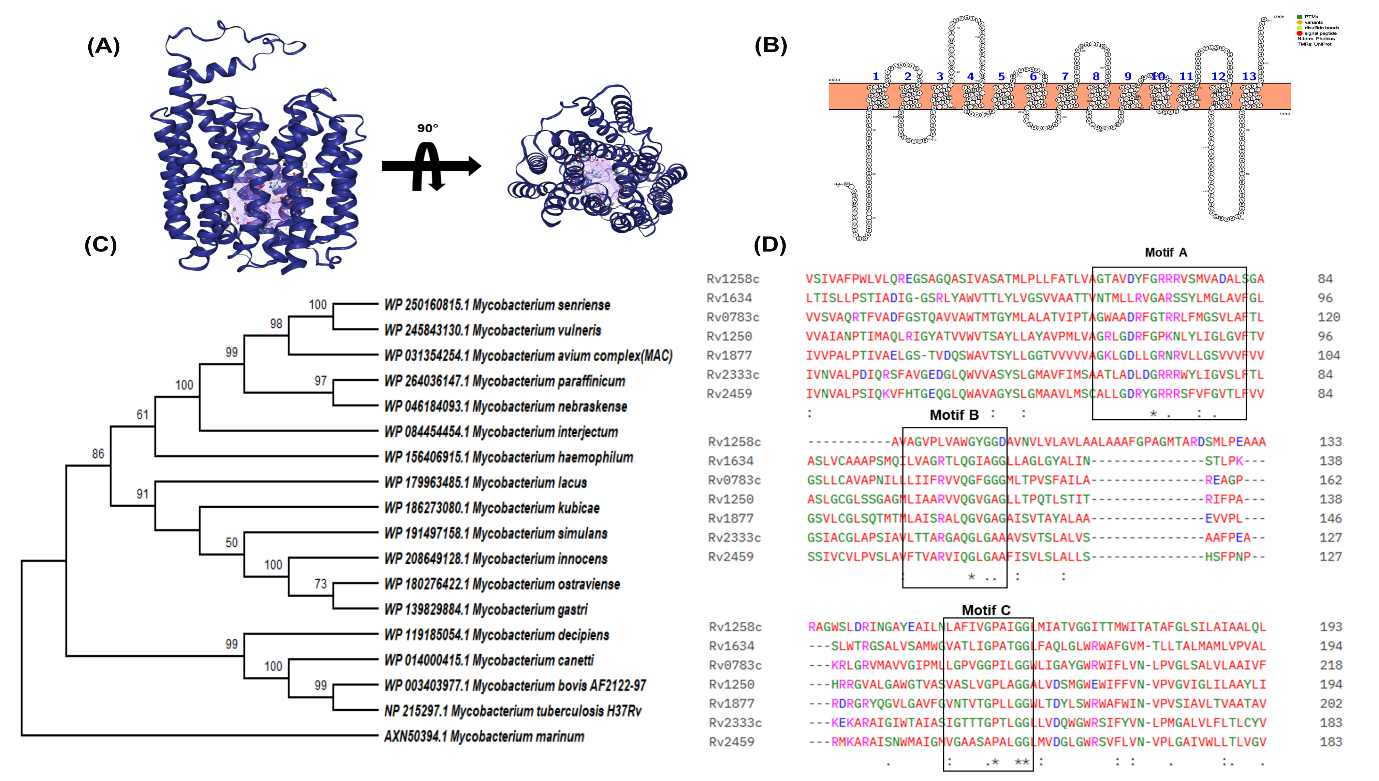

**Figure S1:** **Structural and evolutionary features of Rv0783c.** (A) predicted 3D model from AlphaFold database, with probable pore region viewed by DoGsite scorer server, (B) predicted 13 transmembrane topology showing 13 helices generated by Protter webserver, (C) phylogenetic tree constructed with MEGA12 software showing evolutionary conservation of Rv0783c among different *Mycobacterium species*, (D) multiple sequence alignment (MSA) with MFS transporters identified the conserved motifs-A, B and C.

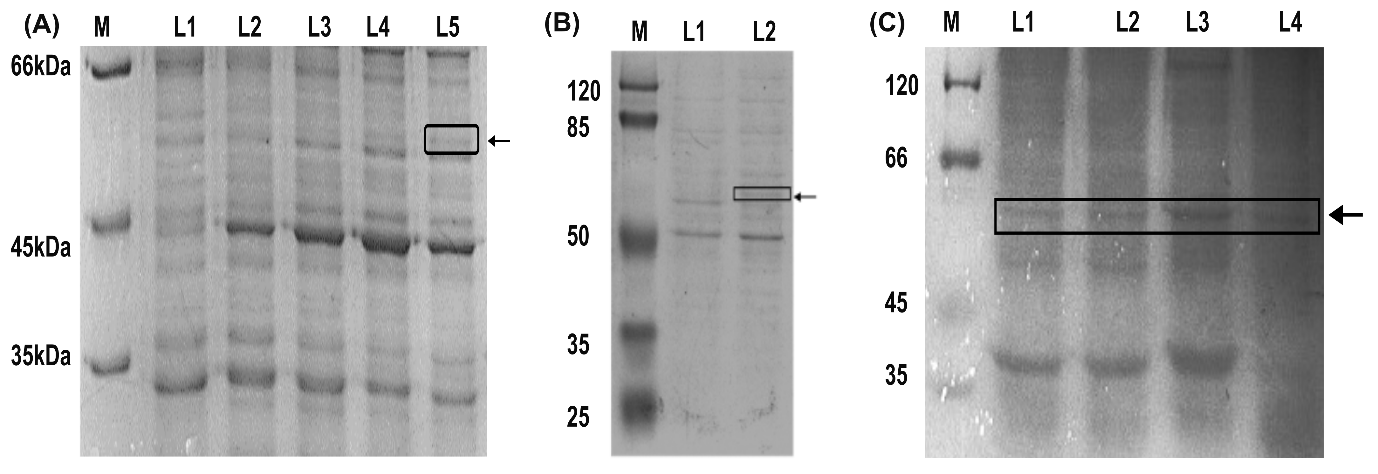

**Figure S2: Expression of Rv0783c from the *E. coli* CS109 and *M. smegmatis*, visualised by 12% SDS-PAGE**. (A) M: unstained molecular weight marker (Thermo Scientific), L1, 2, 3, 4, 5: supernatant fraction from *E. coli* CS109-pBAD, uninduced Rv0783c-pBAD, Rv0783c-pBAD with 0.05%, 0.1%, 0.2% arabinose, respectively, (B) M: prestained molecular weight marker, L1: supernatant fraction of uninduced *M. smegmatis*-Rv0783c, L2: tetracycline-induced *M. smegmatis*-Rv0783c, (C) L1: supernatant fraction of tetracycline-induced *M. smegmatis*-Rv0783c_H56A_, L2: Rv0783c_D58A_, L3: Rv0783c_R138A_, L4: Rv0783c_Q141A._

**Table S1:** Comparative susceptibilities of *E. coli* cells expressing pBAD18-Cm (control and Rv0783c) towards different antimicrobials and compounds, in the absence and presence of sub-inhibitory concentration of CCCP.

| Antibiotic class | Antibiotics | MIC (mg L^-1^) | | | | |
| --- | --- | --- | --- | --- | --- | --- |
|  |  | **Without CCCP** | | | **With CCCP** | |
|  |  | ***E. coli*** | **Rv0783c** |  | ***E. coli*** | **Rv0783c** |
| Fluoroquinolones  (FQ) | **Sparfloxacin** | 0.0125 | 0.05 |  | 0.003 | 0.006 |
|  | **Lomefloxacin** | 0.05 | 0.1 |  | 0.01 | 0.01 |
| Aminoglycosides | **Amikacin** | 4 | 16 |  | 1 | 2 |
|  | **Apramycin** | 8 | 16 |  | 1 | 1 |
|  | **Gentamicin** | 1 | 4 |  | 0.5 | 1 |
|  | **Neomycin** | 2 | 4 |  | 0.25 | 0.25 |
| Beta-lactams | **Ampicillin** | 16 | 64 |  | 2 | 2 |
|  | **Oxacillin** | 8 | 16 |  | 1 | 2 |
| Tetracycline group | **Doxycycline** | 2 | 4 |  | 0.5 | 0.5 |
| Dye | **Ethidium Bromide** | 125 | 250 |  | 15.625 | 31.25 |
|  | **Rhodamine B** | 4 | 8 |  | 1 | 2 |

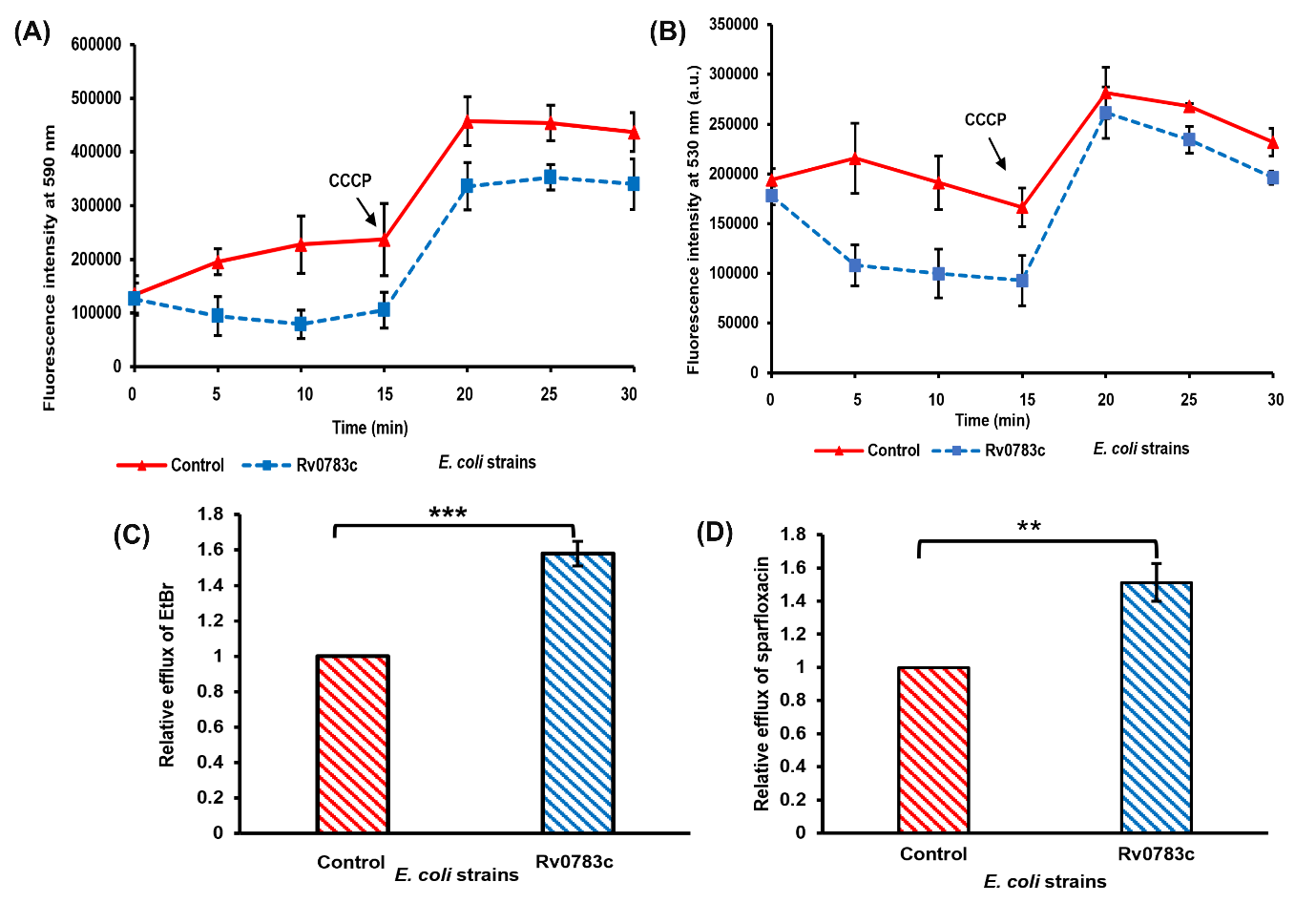

**Figure S3: Intracellular accumulation of antibiotics in *E. coli***. Accumulation of EtBr (A), sparfloxacin (B) at 5 min intervals for 30 min by cells expressing Rv0783c (blue, closed square) compared to the vector control (red, closed triangle). Antibiotics and CCCP were added at 0^th^ and 15^th^ min, respectively. Relative efflux values of sparfloxacin (C), norfloxacin (D) were calculated 10 min after antibiotic addition. Error bars indicate the mean ± standard deviation determined from replicates (n=3). A two-tailed unpaired Student’s t-test was performed with the control and test datasets: ns: non-significant p>0.05, *p<.05, **p<0.01, ***p<0.001, ****p<.0001. The p-values were (C) 0.0015, (D) 0.0001, respectively.

**Table S2:** Comparative docking scores and interacting residues of different antibiotics to Rv0783c, generated by AutoDock-Vina.

| **Antibiotics** | **Binding energy/ Docking score**  **(kcal mol^-1^)** | **Interacting residues** |
| --- | --- | --- |
| Sparfloxacin | -6.2 | H56, T149, Q437, Q436, V59, F314 |
| Lomefloxacin | -6.5 | P180, H56, F314, P184, P321, Q437 |
| Norfloxacin | -7.0 | P180, P184, H56, F317, L318, F317 |
| Ciprofloxacin | -7.2 | H56, P180, P321, F314, V181 |
| Isoniazid | -5.3 | S249, F246, S250, A253 |
| Apramycin | -6.9 | Q437, S249, S441, P252, S448, H56 |
| Amikacin | -6.6 | H56, Q437, F317, M176, G173, F153 |
| Ampicillin | -6.6 | Q436, P180, G173, F314, H56 |

**Table S3:** Comparative susceptibilities of *M. smegmatis* cells with pMIND vector control, Rv0783c and mutated Rv0783c, towards different antimicrobials and compounds.

| **Antibiotics** | **MIC (mg L^-1^)** | | | | |
| --- | --- | --- | --- | --- | --- |
|  | ***M. smegmatis*** | **Rv0783c** | **Q437A** | **G406V** | **F508S** |
| **Sparfloxacin** | 2 | 8 | 8 | 8 | 8 |
| **Lomefloxacin** | 1 | 4 | 4 | 4 | 4 |
| **Norfloxacin** | 2 | 4 | 4 | 4 | 4 |
| **Amikacin** | 2 | 8 | 8 | 8 | 8 |
| **Streptomycin** | 1 | 4 | 4 | 4 | 4 |
| **Apramycin** | 2 | 4 | 4 | 4 | 4 |
| **Gentamicin** | 16 | 32 | 32 | 32 | 32 |
| **Ampicillin** | 64 | 256 | 256 | 256 | 256 |
| **Rifampicin** | 2 | 16 | 16 | 16 | 16 |
| **Isoniazid** | 32 | 128 | 128 | 128 | 128 |
| **Ethambutol** | 0.25 | 0.50 | 0.50 | 0.50 | 0.50 |

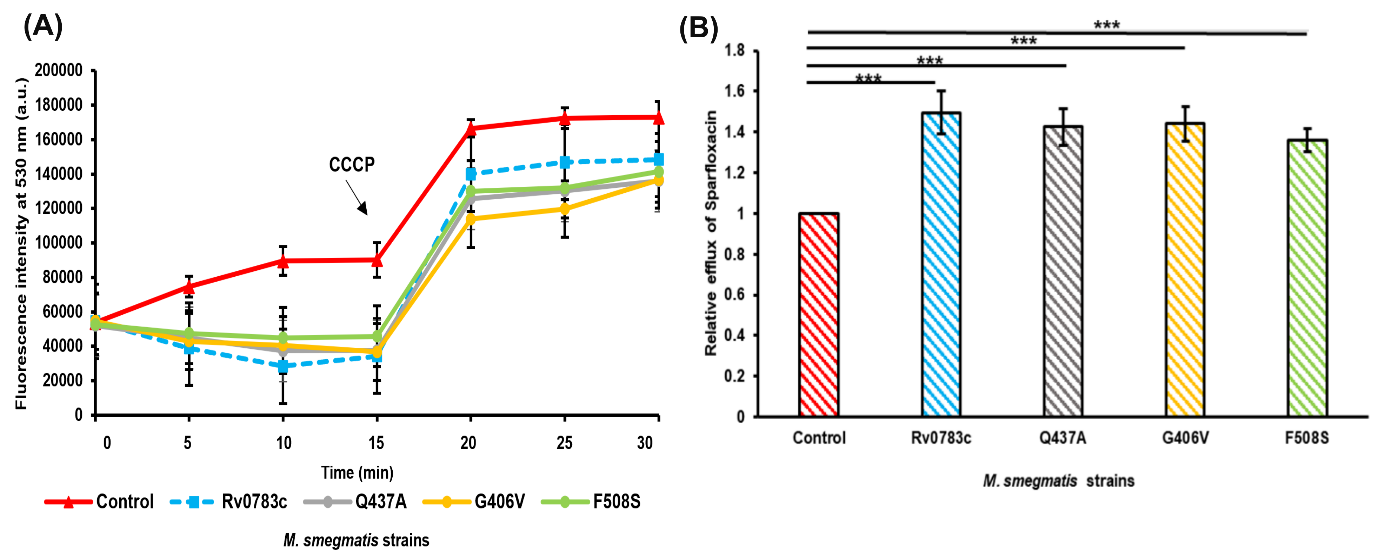

**Figure S4: Sparfloxacin accumulation in *M. smegmatis***. (A) Accumulation of sparfloxacin at 5 min intervals for 30 min by cells expressing Rv0783c (blue, closed square), vector control (red, closed triangle) and mutated Rv0783c. Antibiotics and CCCP were added at 0^th^ and 15^th^ min, respectively. (B) Relative efflux values of sparfloxacin were calculated 10 min after antibiotic addition. Error bars indicate the mean ± standard deviation determined from replicates (n=3). A two-tailed unpaired Student’s t-test was performed with the control and test datasets: ns: non-significant p>0.05, *p<.05, **p<0.01, ***p<0.001, ****p<.0001.

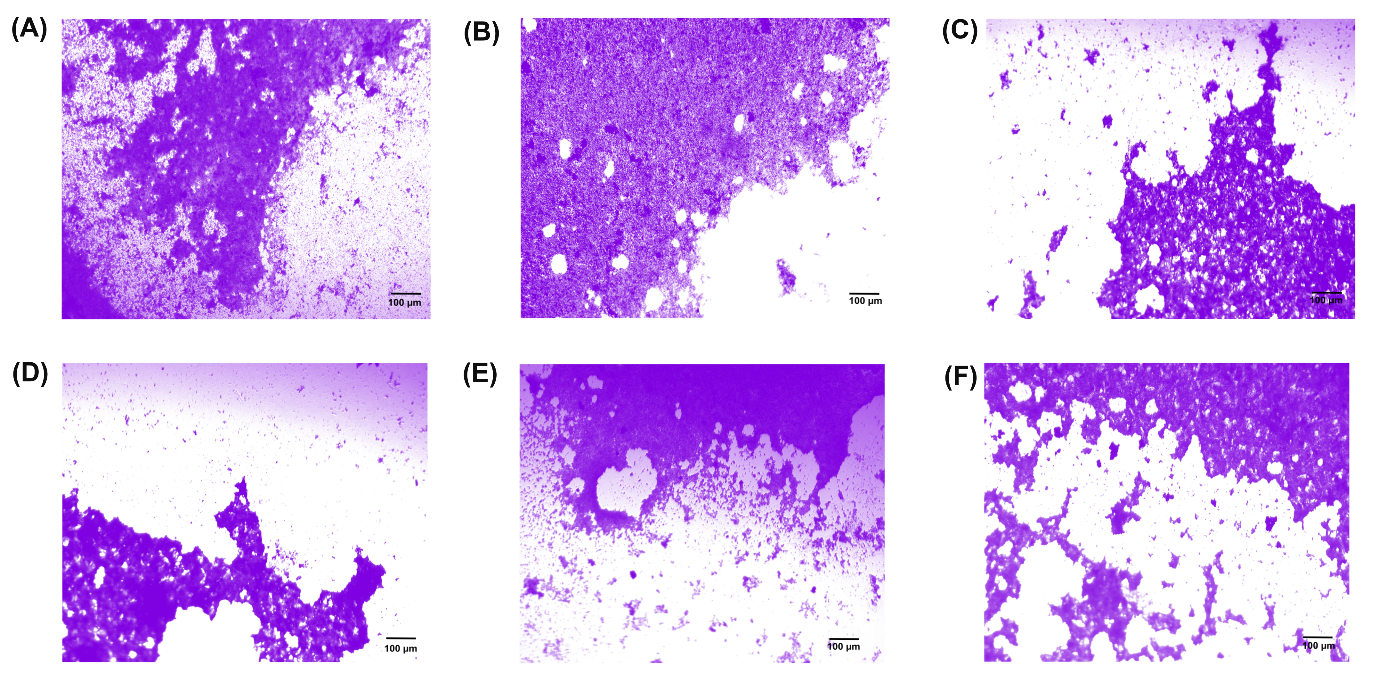

**Figure S5: Microscopic analysis of CV-stained biofilms.** Bright field microscopic images (10×) of the biofilms formed at the air–liquid interface after staining with CV, (A) *M. smegmatis* with pMIND vector, (B) *M. smegmatis* cells expressing Rv0783c_WT_, (C) Rv0783c_H56A_, (D) Rv0783c_D58A_, (E) Rv0783c_R138A_, (F) Rv0783c_Q141A_.

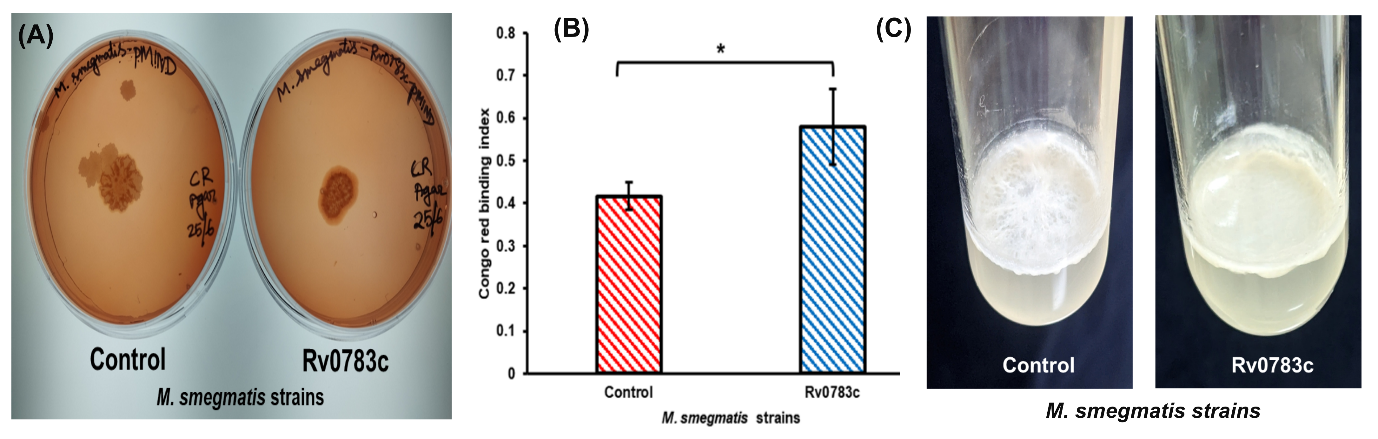

**Figure S6: Congo-red binding assay.** Colony appearance of *M. smegmatis* control and experimental strains on CR supplemented 7H11 agar (A), Congo red binding index derived from OD_488_ of the acetone extracts divided by OD_600_ of the cultures and plotted in a bar graph (B), and pellicle formation on the air-liquid interface after incubation for 7 days (C).

**Table S4:** List of primers used in this study for cloning and site-directed mutagenesis

| **Primer Name** | **Sequence (5’-3’)** |
| --- | --- |
| **pB0783c-FP** | CTCTCTGCTAGCAGGAGGCTCTCTCTATGCTCGGCAACGCCATG |
| **pB0783c-RP** | CTCTCTAAGCTTTCATGCGGATAGCAACGGTGCTCTTCGATGAC |
| **pM0783c-FP** | CTCTCTCATATGAGGAGGCTCTCTCTATGCTCGGCAACGCCATGG |
| **pM0783c-RP** | CTCTCTACTAGTTCATGCGGATAGCAACGGTGCTCTTCGATGAC |
| **0783c_H56A_** | FP: GCCTCGGTGATGGCAGCTGTGGACGTCACCGTG |
|  | RP: CACGGTGACGTCCACAGCTGCCATCACCGAGGC |
| **0783c_Q437A_** | FP: ATCAGCGTCAACCAGGCGGTGGGCGGTTCGATAG |
|  | RP: TATCGAACCGCCCACCGCCTGGTTGACGCTGAT |
| **0783c_F508S_** | FP: GCCTACGCGGTGGTATCCGTGATAGCGACCGCG |
|  | RP: CGCGGTCGCTATCACGGATACCACCGCGTAGGC |
| **0783c_G406V_** | FP: ATGGGCATGGGCATGGTCTGCTCCATGATGCCAC |
|  | RP: TGGCATCATGGAGCAGACCATGCCCATGCCCAT |
| **0783c_D58A_** | FP: GTGATGGCACATGTGGCCGTCACCGTGGTCAGC |
|  | RP: GCTGACCACGGTGACGGCCACATGTGCCATCAC |
| **0783c_R138A_** | FP: CTGCTCATCATATTTGCTGTTGTCCAGGGTTTCGG |
|  | RP: CCGAAACCCTGGACAACAGCAAATATGATGAGCAG |
| **0783c_Q141A_** | FP: ATATTTCGTGTTGTCGCGGGTTTCGGTGGGGGC |
|  | RP: GCCCCCACCGAAACCCGCGACAACACGAAATAT |
